# A *Krüppel-like factor 1* (*KLF1*) mutation associated with severe congenital dyserythropoietic anemia alters its DNA-binding specificity

**DOI:** 10.1101/774158

**Authors:** Klaudia Kulczynska, James J Bieker, Miroslawa Siatecka

**Author notes:** To whom correspondence should be addressed: Miroslawa Siatecka: Department of Genetics, Institute of Experimental Biology, University of Adam Mickiewicz, Poznan 61-614, Poland.

## Abstract

Krüppel-like factor 1 (KLF1/EKLF) is a transcription factor that globally activates genes involved in erythroid cell development. Various mutations are identified in the human KLF1 gene. The E325K mutation causes congenital dyserythropoietic anemia (CDA) type IV, characterized by severe anemia and non-erythroid-related symptoms. The CDA mutation is in the second zinc finger of KLF1 at a position functionally involved in its interactions with DNA. The molecular parameters of how CDA-KLF1 exerts its biological effects have not been addressed. Here, using an in vitro selection strategy we determined the preferred DNA-binding site for CDA-KLF1. Binding to the deduced consensus sequence is supported by in vitro gel shifts and by in vivo functional reporter gene studies. Two significant changes compared to WT binding are observed: G is selected as the middle nucleotide and the 3’-portion of the consensus sequence is more degenerate. As a consequence CDA-KLF1 did not bind the WT consensus sequence. However, activation of ectopic sites is promoted. Continuous activation of WT target genes occurs if they fortuitously contain the novel CDA site nearby. Our findings provide a molecular understanding of how a single mutation in the KLF1 zinc finger exerts an effects on erythroid physiology in CDA type IV.

## Introduction

Krüppel-like Factor 1 (KLF1/EKLF) is an erythroid specific transcription factor essential for red blood cell development (1,2). It has a typical modular structure for transcription factors with its transactivation domain at the N-terminus and three zinc fingers (C2H2) generating its DNA binding domain at the C-terminus. KLF1 belongs to the KLF family of transcription factors that binds the G-rich strand of so-called CACCC-box motifs located in regulatory regions of numerous erythroid genes. KLF1 is involved in activation of globins, globin chaperons, cytoskeleton and membrane proteins, ion and water channels, iron metabolism and heme synthesis enzymes and cell cycle regulators (3,4). Mouse knockout studies have shown that KLF1 ablation is embryonic lethal at day E 14.5 (5,6). KLF1 exerts its function by interacts with chromatin-modifying and -remodeling factors, such as P/CAF, CBP/p300, and the SWI/SNF complex (7-9, reviewed in 10) that help to coordinate opening of chromatin structure, for example at the β-like globin locus (11). Furthermore, KLF1 functions are regulated by various posttranslational modifications such as phosphorylation (12), sumoylation (13), acetylation (7,14), and ubiquitination (15).

The importance of KLF1 for human erythropoiesis is supported by the hematologic diseases and disorders that arise due to mutations occurring along the whole *KLF1* gene. These mutations lead to a range of phenotypic pathology from benign to severe (16,17). Some mutations in KLF1 lead to haploisufficiency that affects expression of certain genes (*Lu/BCAM, Bcl11A, HBD*) that are highly sensitive to KLF1 expression levels. These effects are observed in Hereditary Persistence of Fetal Hemoglobin (HPFH) or the rare In(Lu) blood type (18-20).

The mutations causing the most severe phenotypes affect functionally important amino acids. Two examples of such monoallelic mutations have been described. They substitute the amino acids in the zinc finger DNA binding domain at positions that are involved in direct interactions with regulatory elements of KLF1’s target genes. These mutations alter the properties of the protein such that it acquires novel dominant characteristics (reviewed in: 17,21). They adjust its ability to recognize and bind novel target DNA sequences and lead to serious transcriptional consequences. In both described cases KLF1 mutations are responsible for development of severe anemia. One, found in the mouse and called Nan-KLF1, leads to hemolytic neonatal anemia with Hereditary Spherocytosis (22-24). The second is found in humans and leads to Congenital Dyserythropoietic Anemia type IV (CDA IV) (25,26).

The mouse mutation (E339D), although conservative, on the one hand limits recognition of the normal set of WT-KLF1 targets, but on the other hand leads to acquisition of a novel ability to recognize neomorphic genes not normally activated by WT-KLF1 (24,27,28).

The human substitution is not conservative (E325K), although it is on the same amino acid residue as the mutation of the Nan mice. This charge change on the protein:DNA interface suggests that the consequences for the organism would be much more serious than for the Nan mutation.

To date, only seven patients suffering from CDA type IV have been described (25,26,29-37). In general, CDAs are a heterogeneous group of rare hereditary diseases. The CDA type IV is an autosomal dominant inherited blood disorder (26) characterized by ineffective erythropoiesis and distinct morphological anomalies in the erythroid compartment (blood and bone marrow), such as multinucleated erythroblasts, (38-40), euchromatin areas connecting the nuclear membrane, atypical cytoplasmic inclusions, and intercellular bridges (26,31). In addition, non-erythroid phenotypes such as growth retardation and disturbance in organ development, particularly urogenital anomalies, are manifested in some of these patients (26,29).

In the long run, we are interested in illuminating the molecular mechanism underlying CDA type IV disease and the involvement of mutated CDA-KLF1 in the phenotype. As an essential step towards this ultimate goal we determined the consensus binding site for this mutant. For this purpose we used a CASTing (Cyclic Amplification and Selection of Targets) (41-44) technique, which we combined with next generation sequencing (45). The identified binding motif was verified by gel retardation assays, where we followed complex formation with the newly determined sites. Next, reporter gene assays allowed us to verify transcriptional functionality of these sites.

Our findings reveal that CDA-KLF1 recognizes and binds sites that are mutually exclusive compared to the WT-KLF1 transcription factor. These altered properties have functional consequences that may help explain some of the phenotypic changes seen in the CDA patient’s erythroid cells.

## Results

### Identification of the CDA-KLF1 consensus DNA-binding sequence

To empirically determine the DNA binding consensus site to which CDA-KLF1 binds, first we introduced CDA mutation (E339K) to WT-KLF1, which is an equivalent of E325 position in human. This mutation is found in the second zinc finger (ZnF2) in the “Y” position directly involved in the interactions with DNA (46,47) (Fig.1*A*). Both human and mouse KLF1 orthologues share a very high homology, with 91% identity within the zinc finger domain (Fig. 1*B*). Since most of the KLF1 analyzes were performed on mice, we wanted to change only one variable and follow its effect by direct comparison with the WT and Nan-KLF1 variants. The amino acid located in the “Y” position of ZnF2 interacts with the nucleotide in the middle, 5^th^ position of the 9-nt long binding site (47,48). We started our research by analyzing this relationship. We performed gel retardation assay and followed the complex formation between KLF1 variants which differ in the “Y” position and binding sites with the altered middle 5^th^ position. We examined the binding ability of WT-KLF1 with glutamic acid (E) at position 339 and mutants: CDA with lysine (K) and Nan-KLF1 with aspartic (D) (Fig.1*C*). As binding sites, we used natural KLF1 targets: β-globin with “T” in the middle 5^th^ position, p21 with “C” and E2F2 site 1 with “G” (although WT-KLF1 binds only E2F2-2 and E2F2-3 sites of E2F2 (49)). For “A” in the middle 5^th^ position we have prepared an artificial E2F2-1A. To be sure where the KLF1:double stranded (ds) DNA oligo complex migrates, we added antibodies against KLF1 DNA binding domain epitope, which prevents complex formation (Fig. 1*D*).

**Figure 1.**
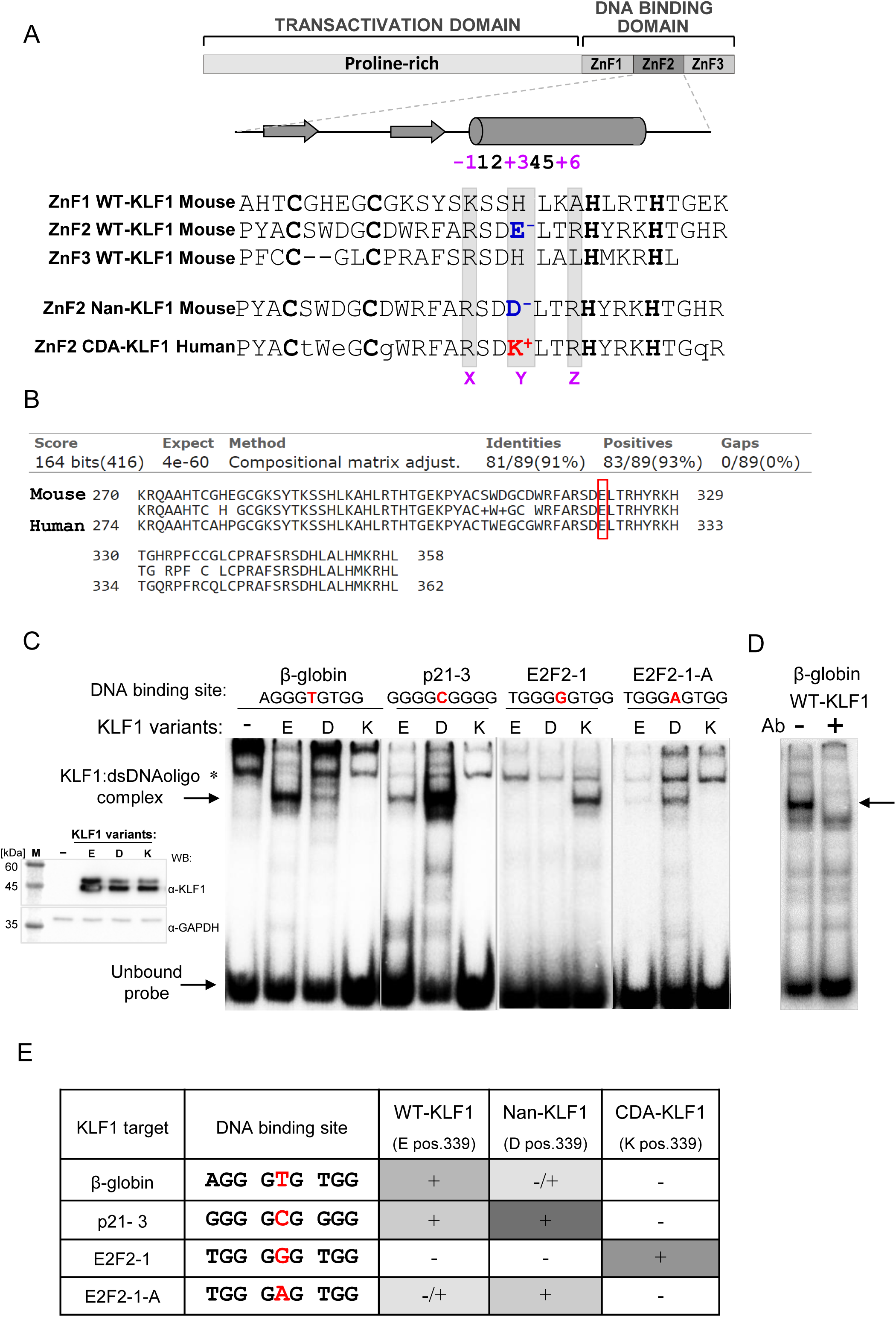
DNA recognition and binding specificity of CDA-KLF1 versus WT-KLF1 and Nan-KLF1. *(A)* A schematic representation of KLF1 consisting of two domains. In the secondary structure of ZnF, arrows indicate β-sheets and the cylinder indicates the α-helix. The alignment of three ZnFs for mouse WT-KLF1, together with mouse ZnF2 of Nan-KLF1 and human ZnF2 of CDA-KLF1, is done based on Cys and His amino acids (bold) involved in zinc coordination. The amino acids denoted X, Y, Z in positions −1, +3, and +6 of the α-helix involved in DNA interactions (46) are shaded in grey boxes. The crucial amino acids in position +3 of ZnF2 are in blue and red according to their charge. *(B)* Amino acid comparison of the murine and human KLF1 proteins (BLASTP). Sequence alignment comprises DNA binding domain, top line represents the mouse KLF1 sequence and the bottom line is the human KLF1. The red box indicates conservative glutamic acid position 339 in mouse and 325 in human. *(C)* Complex formation (shown by arrow) between KLF1 variants (WT -”E”, Nan -”D” and CDA - “K”) and dsDNA oligos with 4 different nucleotides in 5^th^ position (in red). *-nonspecific band. Inset: Western blot with the expression level of KLF1 variants (2µl) used in EMSA (Fig. 1*C*), loading control visualized by antibodies against GAPDH. *(D)* Identification of complex formed between WT-KLF1 and dsDNA β-globin binding site by addition of antibodies (Ab) against KLF1 that prevents complex formation. *(E)* Summary of the interactions between KLF1 variants and the dsDNA oligos based on EMSA (Fig. 1*C*). All possible nucleotides in the 5^th^ position (in red) were tested. The shade of grey represents the intensity of the generated complex.

The results were very exciting (Fig. 1*C, E*), showing that the CDA-KLF1 mutant was able to bind only one motif containing “G” in the middle 5^th^ position. At the same time, two other variants, WT-KLF1 and Nan-KLF1, were unable to bind such a motif, demonstrating the mutually exclusive preferences for binding of the tested proteins. After obtaining these results, we decided to define the binding site for CDA-KLF1 more precisely.

We performed the *in vitro* CASTing-Seq assay, which was combined with high-throughput sequencing to characterize the high-resolution DNA-binding specificity of the KLF1 mutant (Fig. 2*A*). In CASTing we used purified recombinant proteins comprising the DNA binding domain (DBD) of KLF1, which consists of three zinc fingers of the C2H2 type (ZnF-KLF1). The transactivation domain contains numerous prolines that make the structure unstable and prone to degradation, rendering it impossible to use the full length protein in this assay. The DBD domain had His- and Flag-tags fused at the N-terminus, enabling its purification by His-tag affinity followed by removal of the His-tag. The resultant recombinant protein was bound via its Flag-tag to magnetic beads coated with Anti-Flag M2 monoclonal antibody and used in CASTing assay. In parallel to CDA-ZnF-KLF1, we used wild type DBD (WT-ZnF-KLF1) as a positive control, for which the consensus binding site is already known (4). The magnetic beads alone without any protein attached served as a negative control. At first we used the library that contained 12 random nucleotides flanked by constant sequences needed for amplification. The magnetic properties of the beads allowed for a very rapid separation of the beads coated by protein with bound DNA oligonucleotides from a suspension containing unbound oligonucleotides. The remaining DNA oligonucleotides that interacted with the particular ZnF domains were PCR amplified, purified and used for the next round of the assay. We performed five such cycles.

**Figure 2.**
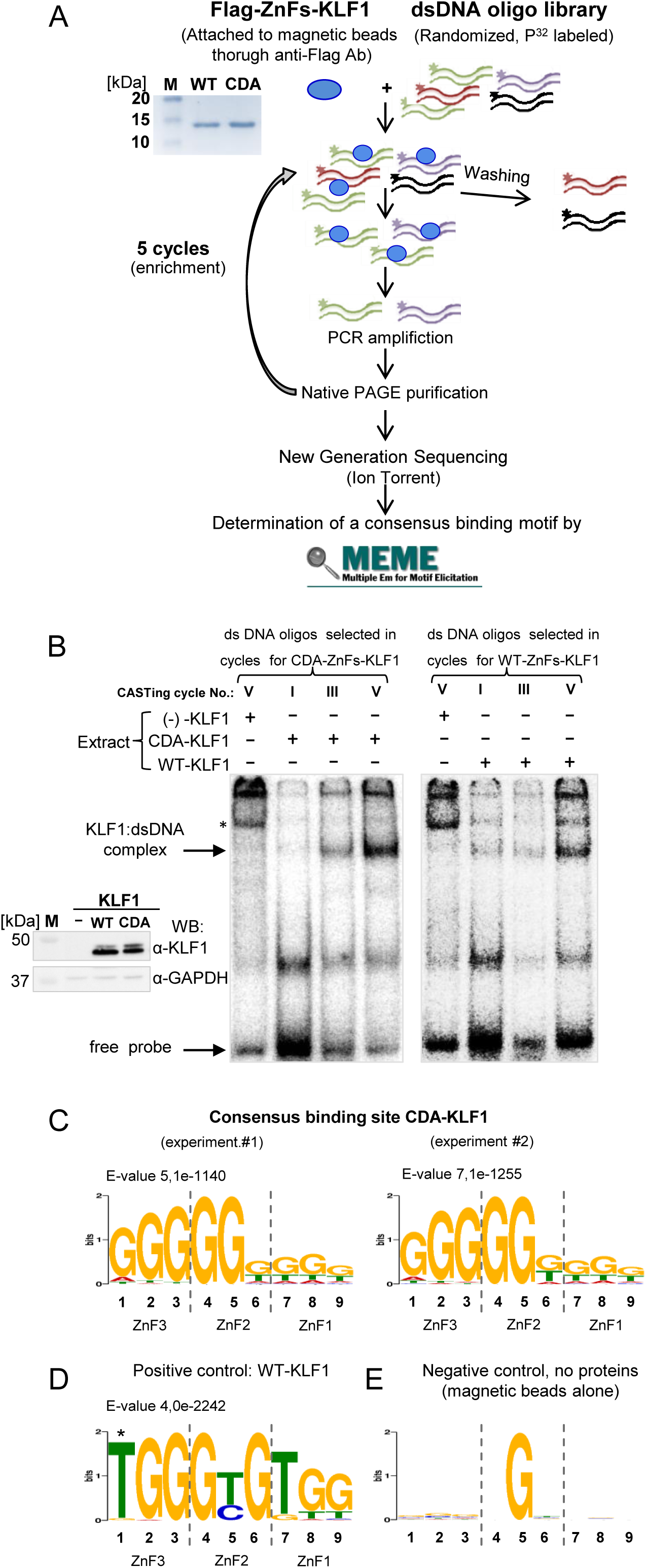
Determination of a consensus binding sites for CDA variant of KLF1 using the CASTing-Seq technique and MEME motif-based sequence analysis tools. *(A)* The scheme of CASTing-Seq technique is shown, from protein/library incubation to cycling and determination of the consensus binding motif. The inset shows Coomassie stained purified recombinant proteins that were the starting material and comprise the three zinc finger domains of WT-KLF1 and CDA-KLF1, respectively. *(B)* Enrichment of oligonucleotide pools selected in CASTing cycles I, III and V for ZnF-KLF1 variants. Gel shift assay analyses showing an increased efficiency in complex formation (shown by arrow) between radiolabeled dsDNA oligonucleotides from cycle number: I, III and V of CASTing and full length CDA-KLF1 (left panel) and WT-KLF1 (right panel) in extracts from transfected COS-7 cells. * -nonspecific band. The inset shows western blot of extracts used for the gel shift analysis. *(C)* A new consensus DNA binding motif for CDA-ZnF-KLF1 obtained in two biological repeats of CASTing-Seq experiment and MEME motif-based sequence analysis. *(D)* The DNA binding motif obtained for WT-ZnF-KLF1 in CASTing-Seq experiment that serves as a positive control. *Star* marks thymidine that comes from the flanking sequence of the random core of the oligonucleotide library. *(E)* Lack of selection of any DNA binding motif for the magnetic beads alone that serves as a negative control in CASTing-Seq experiment.

The DNA oligonucleotides enriched after fifth cycle were sequenced by NGS Ion Torrent. The results showed that CDA-ZnF-KLF1 recognizes a very G-rich sequence, containing guanine in the middle (5^th^) position of the 9-nt binding motif (data not shown). This nucleotide is directly involved in interactions with the amino acid in position 339 of the second zinc finger of KLF1 (24,50). In the wild-type scenario, WT-KLF1, with glutamic acid (E) in position 339, recognizes a 5’-NGG-GC/TG-T/GGG-3’ consensus site; that is, a T or C in the middle position. Thus, oligonucleotides containing G in the 5^th^ position were not expected to form a complex with WT-KLF1 (24). Our results from CASTing-Seq assay suggested that the CDA mutation (E339K) alters the binding specificity of KLF1. It was in agreement with our initial observation, that CDA-KLF1 recognizes G in the middle position of binding site (Fig. 1*C*). However, because of the long stretches of G repeats found in the CASTing-selected CDA-ZnF-KLF1 binding motifs, and due to Ion Torrent system limitations related to sequencing of homopolymer repeats, it was difficult to unambiguously determine the CDA-KLF1 consensus binding motif.

In order to overcome this problem we repeated the CASTing assay, but this time we applied a more accurate library. In this approach, flanking sequences surround 9-nt long DNA oligonucleotides 5’- NNN-N<u>G</u>N-NNN-3’, which are precisely the length of binding site for three zinc fingers. In addition, we forced G in the middle position, leaving the rest random and the entire CASTing-Seq experiment was performed once again. The gradual enrichment of selected oligonucleotides from the library in cycles I, III and V DNA was monitored by EMSA assay. The EMSA complexes were formed with full length CDA-KLF1 or WT-KLF1, expressed in extracts of transfected COS-7 cells (Fig. 2*B*), which therefore verified that the CASTing oligonucleotides selected by the zinc finger domain interact with the full length proteins.

The DNA oligonucleotides obtained after final round were sequenced by Ion Torrent. The resultant ∼ 30 000 sequences were analyzed on a Galaxy platform and then the MEME suite was applied. The final results are summarized on Fig. 2*C*-*E*. The new consensus binding site for CDA-KLF1 was determined to be 5’-NGG-G<u>G</u>T/G-T/GT/GT/G-3’. The data for CDA-ZnF-KLF1 were obtained in two biological repeats (Fig. 2*C*). The reliability of the results is based on the positive control data (WT-ZnF-KLF1; Fig. 2*D*), which are in agreement with published consensus site 5’-NGG-GC/TG-T/GGG-3’ (4,24,27). The thymine in first position marked with a star (Fig. 2*D*) comes from flanking sequence of the library. By way of explanation, the second library of the random oligonucleotides with G fixed in the middle (5^th^) position of the motif was designed especially for CDA-ZnF-KLF1. Thus in order to be bound by WT-ZnF-KLF1, a shift has to occur to have C or T in the middle position. In such a situation the binding motif had to expand to the flanking sequence. Indeed, the consensus sequence motif for WT zinc finger domain contains C or T in the 5^th^ position as expected and a flanking T in the 1^st^ position. The negative control, the magnetic beads alone, did not select any sequence motif; it remained random (Fig. 2*E*).

In conclusion, we have newly identified the CDA-KLF1 binding motif. It is G-rich, it has G in the middle (5^th^) position crucial for interactions with ZnF2 pos.339 of KLF1 and it shows rather high degeneracy at the 3’ end of the motif. The degeneracy refers to the positions 6^th^, 7^th^, 8^th^ and 9^th^ on the G-rich strand that are mostly involved in interactions with ZnF1 (Fig.2*C*).

Next, we validated the *in vitro* selection results by EMSA assays, functional reporter assays in K562 cells and *in vivo* analyses using cell line based KLF1 variant overexpression assay. Several important issues were analyzed in this set of experiments: (1) confirming the specificity of KLF1 variants based on the 5^th^middle position of binding site (2) testing individual oligo sequences with their 3’ degeneracy and their effect on interactions with ZnF domains; (3) testing the specificity of the interactions between KLF1 variants and particular oligonucleotides and (4) *in vivo* analysis of expression level of selected KLF1 targets upon induction of CDA or WT-KLF1.

### Analysis of the essential G in the middle (5^th^) position of binding site for CDA-KLF1

We started validating the *in vitro* consensus binding site for the KLF1 CDA mutant by performing a quantitative EMSA experiment and a dissociation constant measurement. The complexes were generated with an increased amount of purified, recombinant zinc finger domain for WT or CDA-KLF1, respectively (Fig. 3). We monitored the formation of complexes for one of the newly determined 5’-GGG-G<u>G</u>G-GGG-3’ site and for the β-globin binding site 5’-AGG-G<u>T</u>G-TGG-3’, as a positive control for WT-KLF1. We measured dissociation constant (K_d_) for all the possible combinations of complexes (Fig. 3).

**Figure 3.**
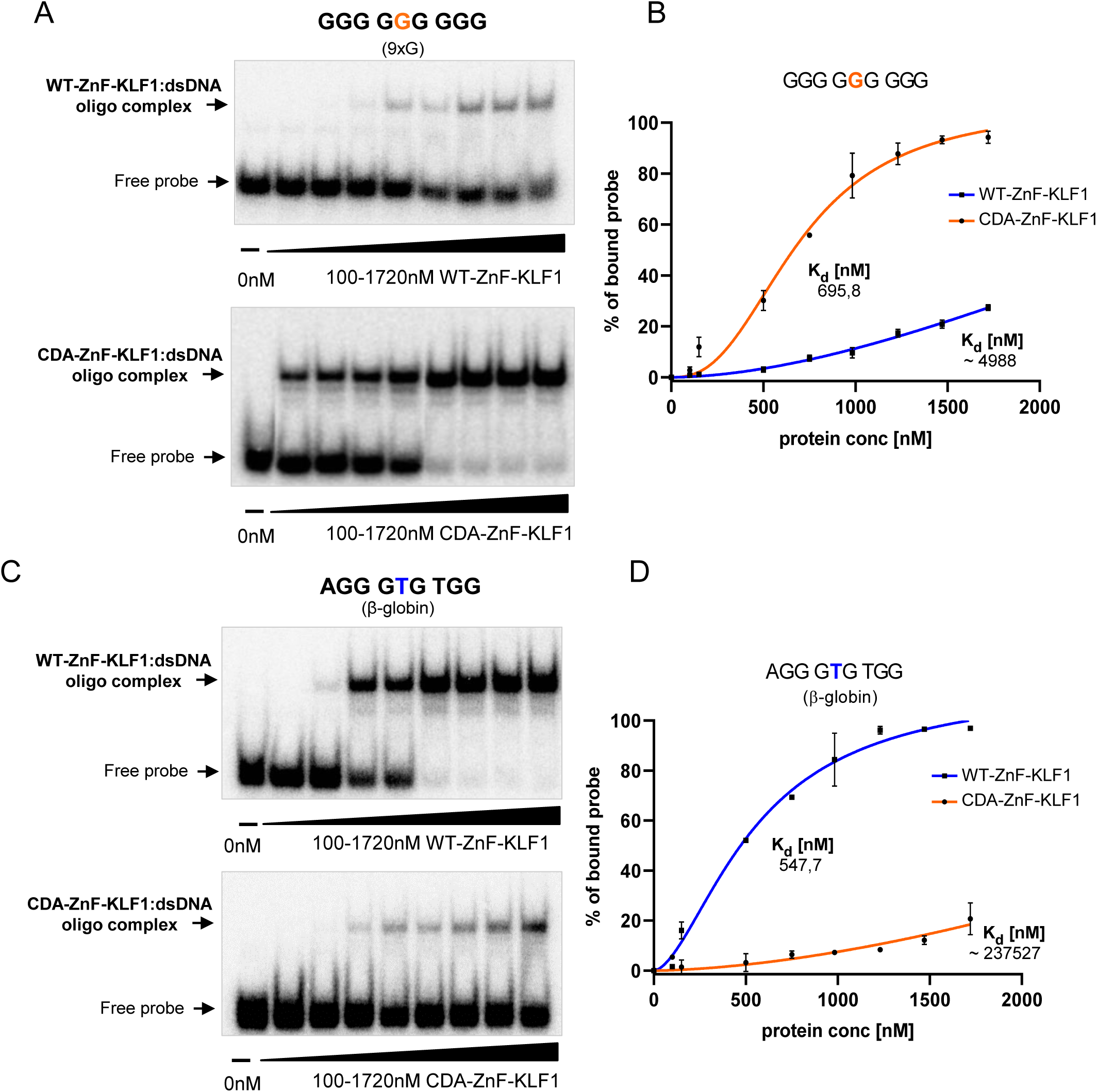
Comparison of the binding affinity of the zinc finger (ZnF) domain of WT-KLF1 and CDA-KLF1 towards newly identified binding site with G in the middle 5^th^ position (9xG) and β-globin binding site. *(A)* Complex formation between an increased amount (0-1720nM) of purified recombinant ZnF domain (WT or CDA) and radiolabeled dsDNA 5’-GGG-G<u>G</u>G-GGG-3’ binding site newly identified for CDA-KLF1. *(B)* K_d_ binding curves for 9xG dsDNA oligo and KLF1 variants as indicated, obtained using Hill Slope in GraphPad Prism. Average K_d_ were calculated based on at least two EMSA gels. *(C)* Complex formation between an increased amount (0-1720nM) of purified recombinant ZnF domain (WT or CDA) and radiolabeled dsDNA 5’-AGG-G<u>T</u>G-TGG-3’ binding site for β-globin. *(D)* K_d_ binding curves for β-globin dsDNA oligo and KLF1 variants as indicated obtained using Hill Slope in GraphPad Prism. Average Kd were calculated based on at least two EMSA gels.

Under applied conditions the binding affinity of WT-ZnF-KLF1 to β-globin site is K_d_= 547,7 ± 76,4nM and it is approximately 10-fold lower than for binding site with G in the middle position (estimated K_d_ ∼ 4988nM). The situation was reversed for CDA-ZnF-KLF1, which binds 5’-GGG-GGG-GGG-3’ with an affinity of K_d_=695,8 ± 71,1nM, while the dissociation constant for the β-globin binding site increased dramatically to K_d_ approx.∼ 237527nM (Fig. 3*A-D*). These results clearly indicate preferences in the specificity of KLF1 variants based on the middle 5^th^ position of the binding site.

Next, we performed a cold competition assay which additionally supports the specificity of CDA-KLF1 binding to a new site with G in the middle position (Fig. 4). For this purpose we used full-length WT or CDA-KLF1 protein from transfected COS-7 extracts and performed EMSA. Complexes between CDA-KLF1 and radiolabeled dsDNA oligo with G in the middle 5^th^ position were effectively antagonized by cold competition with binding sites that contain middle G and are marked as E2F2-1 and 9xG (Fig. 4 panel A, B). In the case of WT-KLF1 complexes with radiolabeled β-globin binding site, they were replaced only with cold β-globin sites. Two other binding sites with G in the middle 5^th^ position, i.e. E2F2-1 and 9xG did not affect the WT-KLF1 complexes (Fig.4 panel *C*).

**Figure 4.**
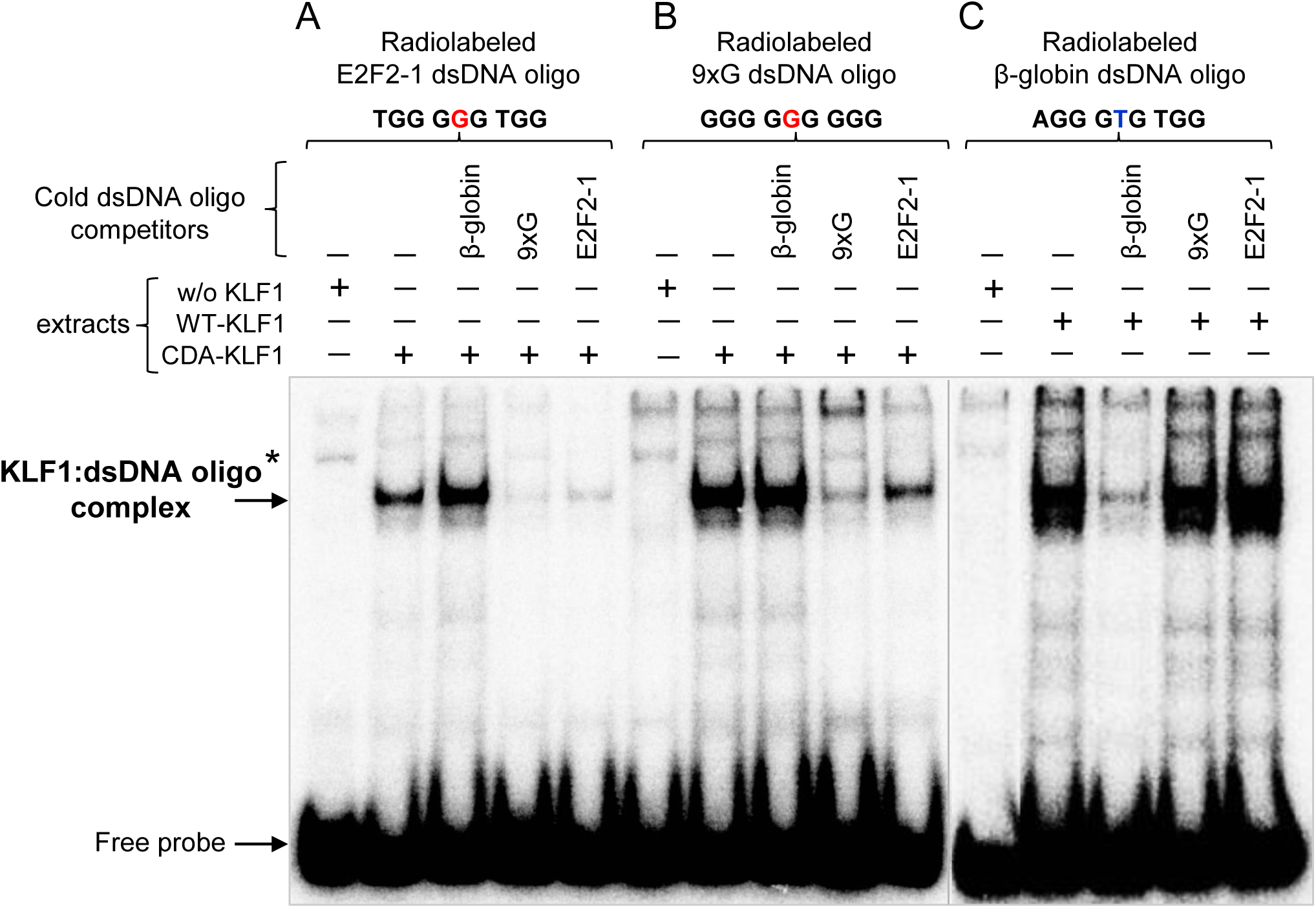
CDA-KLF1 binding specificity of the newly determined consensus binding site. Gel shift assay monitoring complexes formation (indicated by arrow) between full length CDA-KLF1 (part A and B) with radiolabeled dsDNA oligonucleotides: E2F2-1 (5’-TGG-G<u>G</u>G-TGG-3’) and 9xG (5’-GGG-G<u>G</u>G-GGG-3’), respectively and WT-KLF1 (part C) with radiolabeled dsDNA β-globin (5’-AGG-G<u>T</u>G-TGG-3’) binding site. Complexes formed by both KLF1 variants were subjected to a binding competition with a 100-fold molar excess of the indicated unlabeled (cold) ds oligonucleotides. *-nonspecific band. The protein extracts used in these EMSAs are the same as visualized on western blot –Fig. 1C.

Both experiments indicate without a doubt that CDA-KLF1 and WT-KLF1 have mutually exclusive binding sites.

### Validation of the CASTing-Seq results focusing on 3’ end degeneracy of the consensus binding site

Several individual CASTing-based sequences were tested in parallel for their specificity in complex formation with CDA (E339K mutation) and with WT (E339) zinc fingers (Fig. 5). We focused on testing of the 3’ degeneracy of the consensus binding site and analyzed oligonucleotides differed in three nucleotides (positions 7^th^, 8^th^ and 9^th^ on the G-rich strand) that are recognized and bound by zinc finger Based on EMSA results we distinguished two major groups of binding sites depending on their ability of complex formation. First, the largest group consists of sites that are bound by CDA-ZnF-KLF1 and not bound by WT-Zn-KLF1 (Fig. 5*A*), and a second group of sites that are not bound by either CDA nor WT-ZnF-KLF1 (Fig. 5*B*). As a positive control for WT-ZnF-KLF1 we used β-globin binding site, which has “T” in the middle 5^th^ position, which at the same time is not a proper site for CDA-ZnF-KLF1 (Fig. 5*C*). The results were subjected to densitometric analysis to estimate the percentage involvement of the tested oligonucleotides in interactions with the CDA or WT-ZnF recombinant domain, respectively.

**Figure 5.**
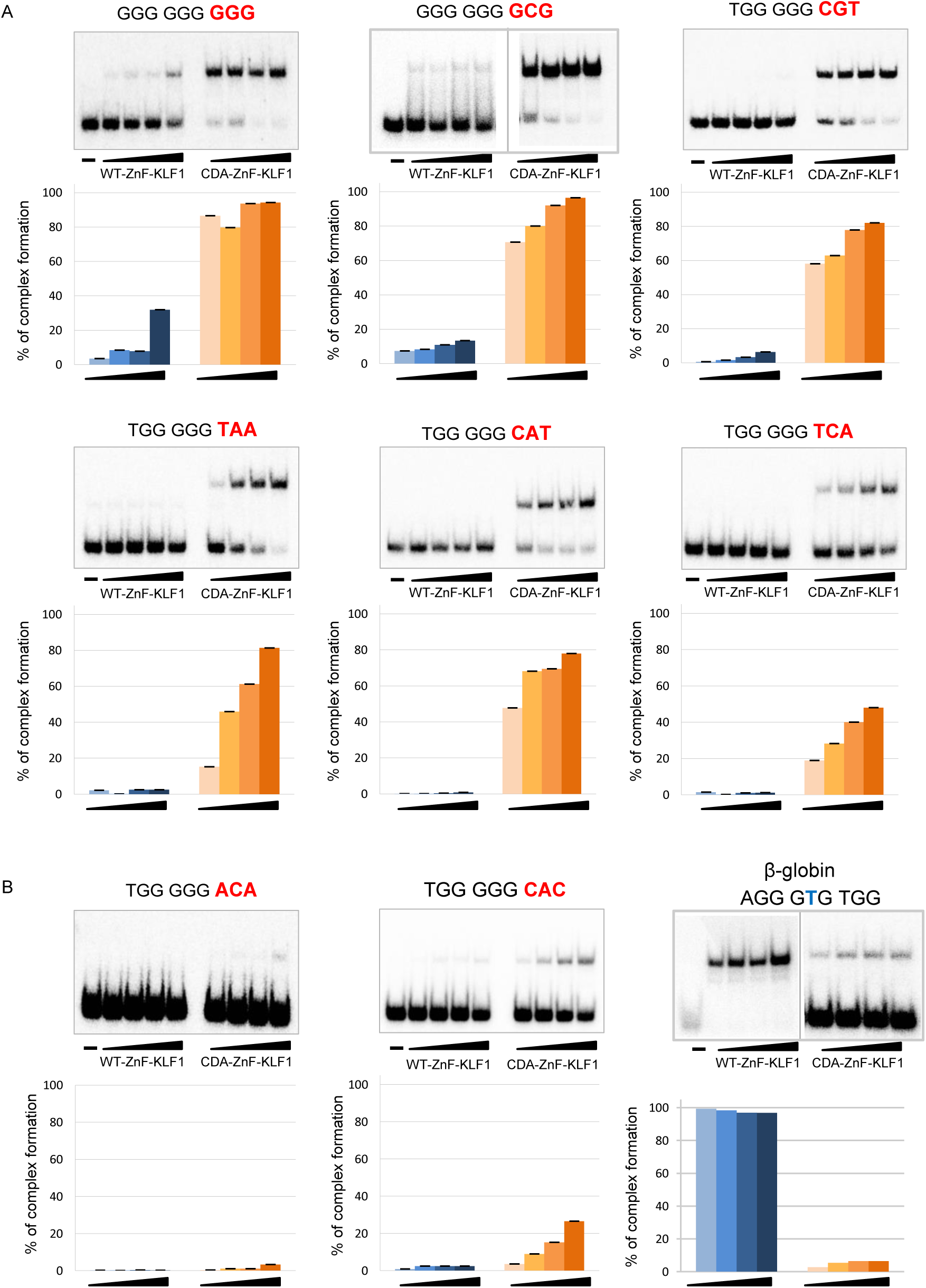
Comparison of the efficiency of complex formation between individual sequences derived from the consensus binding site for CDA-KLF1, and the ZnF domain of WT-KLF1 and CDA-KLF1, respectively. *Upper panel:* Gel shift assay performed with an increased amount (0.12µg, 0.15µg, 0.18µg and 0.21µg) of purified recombinant ZnF domain (WT or CDA) and radiolabeled dsDNA oligonucleotides comprising binding sites for CDA-KLF1. Analyzed DNA sequences are denoted above each individual gel. *Lower panel:* the densitometric analysis of the gel shift results showing the percentage of dsDNA oligo that is involved in complex formation with WT-ZnF-KLF1 and CDA-ZnF-KLF1, respectively. The graphs are made based on a single representative EMSA gels above, but gel shift analyses were performed at least twice for each tested binding site. (*A*) The DNA binding sites that generate efficient complexes with CDA-ZnF-KLF1 as compared to WT-ZnF-KLF1. (*B*) The DNA binding sites that generate very weak complexes with CDA-ZnF-KLF1 comparable to WT-ZnF-KLF1. (*C*) The DNA binding site for β-globin that generates the complex with WT-ZnF-KLF1 domain serves as a positive control.

We find that oligonucleotides having at least one G or T in the 3’ triple nucleotide are efficiently bound by CDA-ZnF-KLF1, whereas complexes generated with oligonucleotides containing only C or A in positions 7^th^ −9^th^ are very weak, estimated below 20% (Fig. 5*B*). The K_d_ values for the complexes support the CDA-KLF1 particular binding preferences as compared to WT-KLF1.

### Analysis of the specificity of complex formation between full length of KLF1 variants (WT, CDA, and Nan-KLF1) and radiolabeled oligonucleotides

As we could use only the DNA binding domain of KLF1 for the CASTing experiment, subsequently, we tested whether full length CDA-KLF1 is able to interact with the determined consensus binding sites. For this purpose we chose the same individual binding sites as previously tested for the 3’ degeneracy (Fig. 5) and performed EMSA assays using extracts from transfected COS-7 cells expressing equivalent amounts of full length constructs (Fig. 6).

**Figure 6.**
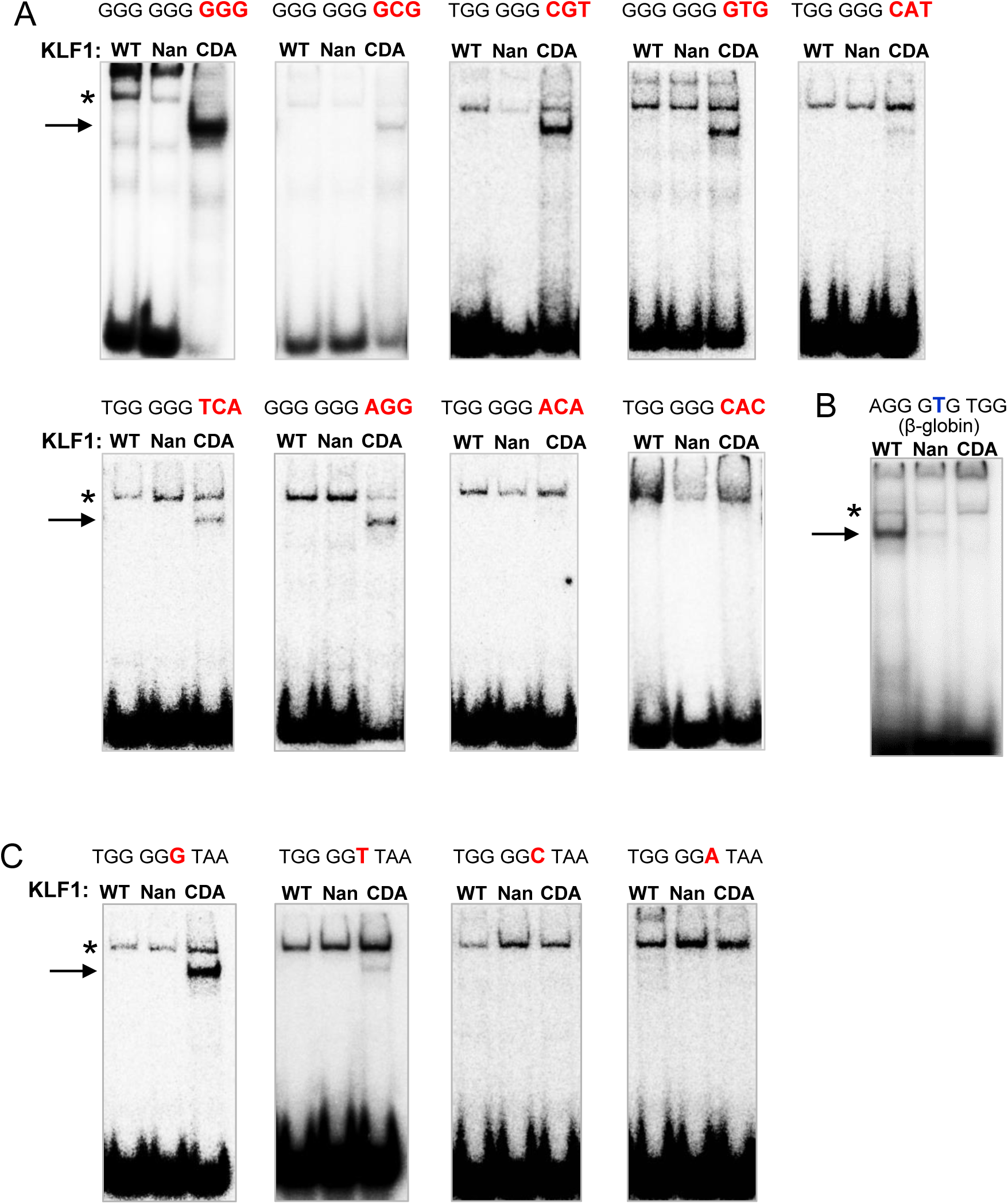
Analysis of the new consensus DNA binding site for CDA-KLF1 and investigation of the binding specificity by KLF1 variants (CDA versus WT and Nan) with an emphasis of analysis of the degenerate 3’ end of CDA-KLF1 consensus binding site. (*A*) Analysis of variations in positions 7^th^, 8^th^, and 9^th^ within the CDA-KLF1 binding site that interact with zinc finger 1 of KLF1. Gel shift assays comparing complex formation (indicated by arrow) between full length WT-KLF1, Nan-KLF1 and CDA-KLF1 from extracts of transfected COS-7 cells and radiolabeled dsDNA oligo binding sites as indicated. *-nonspecific band. (*B*) Complex formation with β-globin binding site serves as a positive control for WT-KLF1. (*C*) Comparable analysis of the degenerate position 6^th^ of the consensus binding site for CDA-KLF1. Four binding sites that differ in position 6^th^ were subjected to complex formation with KLF1 variants, indicated by the arrow, *- nonspecific band. The protein extracts used in these EMSAs are the same as visualized on western blot –Fig. 1C.

Here, in parallel, we tested the specificity of recognition of these oligonucleotides by different KLF1 variants that vary in the amino acid in position 339: WT-KLF1 with E339, Nan-KLF1 with D339, and CDA-KLF1 with K339.

The oligonucleotides that have G in the middle (5^th^) position and the 3’ triple nucleotide consisting of at least one T or G were bound by CDA-KLF1 (Fig. 6*A*) similarly to the zinc finger domain alone (compare Fig. 6*A* and 5*A*). However, the oligonucleotides that have G in middle (5^th^) position of the motif but do not have any G or T in the last triple nucleotide were not bound by CDA-KLF1 (compare Fig. 6*A* and 5*B*). Neither WT-KLF1 nor Nan-KLF1 full length proteins are able to interact with binding sites containing G in the middle position on the G-rich strand in any case, which is consistent with our previous observations (24). As a positive control for these analyses we took again β-globin binding site, which was recognize and bound by WT-KLF1 and slightly by Nan-KLF1 (Fig. 6*B*).

We conclude that the novel binding properties determined by *in vitro* selection with the CDA-KLF1 DBD alone are retained when tested using the full length CDA-KLF1 protein.

Next, we focused on the degenerate 6^th^ position of novel CDA-KLF1 consensus binding site (Fig. 2*C*). We applied gel retardation testing and monitored complex formation between KLF1 variants and radiolabeled oligonucleotides that differ in this 6^th^ position. We analyzed all four combinations of nucleotides and the data reveled that only G or (more weakly) T in position 6^th^ create a suitable binding motif for CDA-KLF1 (Fig. 6*C*). As expected, WT-KLF1 or Nan-KLF1 did not bind to any of these, as they retain G in the 5^th^ position that prevents the interactions.

These results clearly demonstrate that the newly determined CDA-KLF1 consensus binding site, 5’-NGG-G<u>G</u>T/G-T/GT/GT/G-3’, crucially prefers G in the middle (5^th^) position that directly interacts with the mutated E339K residue in the CDA-KLF1 protein. In addition, this single amino acid change yields a protein with an unexpectedly high level of binding degeneracy for the sequence that interacts with zinc finger 1 of CDA-KLF1 (positions 7^th^, 8^th^ and 9^th^ on G-rich strand), as well as in position 6^th^. Thus, CDA-KLF1 preferentially binds sites containing G or T in the last four nucleotides of the consensus binding motif.

### Functional analysis of individual CDA-KLF1 binding sites by luciferase reporter gene assays in K562 cells

To test if the newly identified consensus site for CDA-KLF1 is functional, we used a luciferase reporter gene system that provides a robust and specific readout of KLF1 activity (51). We constructed a set of luciferase reporters where, upstream of the luciferase coding sequence with its minimal TK (thymidine kinase) promoter, we placed 6x repeated individual 9-nt long binding sites selected from CASTing (Fig. 7, top). We used them for co-transfection of K562 cells together with particular KLF1 variants (WT, Nan, or CDA). In our K562 cells the endogenous KLF1 is not expressed (52) thus we can easily investigate the individual activity of the KLF1 variants by measuring the activation of luciferase and monitoring the relative light units (RLUs).

**Figure 7.**
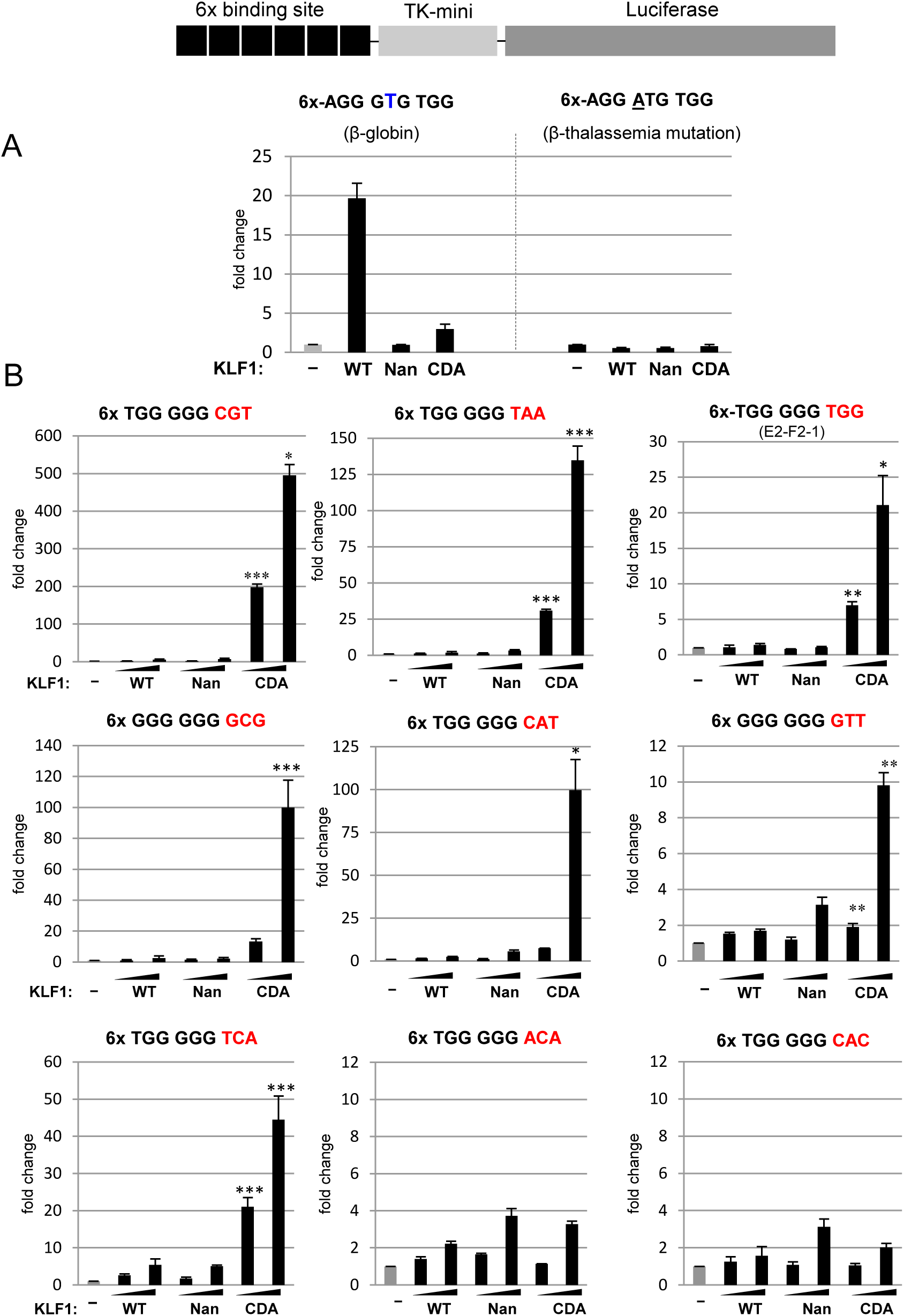
Functional analyses of the specificity of KLF1 variants in recognition and binding of CASTing-selected sites for CDA-KLF1 by activation of luciferase reporter. K562 erythroleukemia cells that do not express endogenous KLF1 (52) were cotransfected with plasmids expressing the luciferase reporter gene along with increasing amounts of WT-KLF1, Nan-KLF1, or CDA-KLF1. *Top:* A scheme of the luciferase reporter gene under control of the 63-bp fragment of *thymidine kinase* promoter (TK mini) and 6x repeated selected binding sites (*A*) Comparison of the fold change of transcriptional ability of full length WT-KLF1, Nan-KLF1, and CDA-KLF1 in binding and activation of luciferase reporter gene with binding site from β-globin promoter (1) served as a positive control and binding site with thalassemia mutation in β-globin promoter (underlined) (53) as a negative control. (*B*) Comparison of the fold change of transcriptional ability of full length WT-KLF1, Nan-KLF1, and CDA-KLF1 in binding and activation a set of luciferase reporter genes that were constructed based on CASTing-selected binding sites as shown at the top of each panel. A *Renilla* reporter construct was included as a normalization control for transfection efficiency. An average of three biological replicates for each reporter gene (arithmetic mean ± SD) is shown. *p≤0.05, ** p≤0.01, ***p≤ 0.001.

Our positive control, which is a reporter containing 6x β-globin binding site 5’-AGG-GTG-TGG-3’, is solely activated by WT-KLF1 but not by Nan- or CDA-KLF1, as expected (Fig. 7*A*). The negative control, which consists of the same 6x repeated β-globin site but with a thalassemia point mutation 5’-AGG-<u>A</u>TG-TGG-3’ (53), is not activated by any KLF1 construct (Fig. 7*A*). These data demonstrate that our assay system is robust and selective.

For the functional analyses we used the same binding site sequences as for *in vitro* EMSA assays described above (Fig. 5 and 6). To make the results more convincing, we transfected K562 cells with increasing amounts of expression plasmids carrying KLF1 variants. We find that the luciferase reporter genes are activated only from binding sites that are recognized and bound by CDA-KLF1 as proven by gel retardation assay (compare Fig. 6*A* with 7*B*). Such sites need to have G in the middle (5^th^) position of the binding motif, and the 3’ end of the sequence (position 7^th^, 8^th^ and 9^th^) has to contain at least one G or T. Neither WT-KLF1 nor Nan-KLF1 are able to activate any of studied reporter genes, because of the G in the middle position.

These results evidently show that only CDA-KLF1 is able to activate the reporter genes in a pattern totally consistent with the previous analyses and thus are entirely confirmatory with the *in vitro* EMSA results.

### Quantitative analysis of the endogenous levels of KLF1 target genes, expressed in stable K562 cell lines with induced expression of WT- or CDA-KLF1

Next, we studied the expression level of endogenous genes that might be affected by the presence of mutant KLF1. For this purpose we prepared two stable K562 cell lines with doxycycline inducible KLF1 variants, either WT-KLF1 or CDA-KLF1. After induction, we isolated total RNA and prepared reverse transcribed cDNA for quantitative analysis. We focused on monitoring expression of known WT-KLF1 targets, notably: *Lu*/*BCAM*-basal cell adhesion molecule (Lutheran blood group), *AQP1*- Aquaporin 1, *EPB4.2* - erythrocyte protein band 4.2, *Cdkn1B*-Cyclin Dependent Kinase Inhibitor 1B (p27), *E2F2*-E2F transcription factor. Our rationale was as follows: first, it is known that AQP1 levels are dramatically lower in samples originating from patients suffering from CDA type IV compared to healthy controls (16,35,54). Expression of *EPB4.2* is reduced in CDA-iPS-derived erythroblasts (37) also analysis of transcriptome of the CDA patient revealed it decreased level, as it contributes to the fragility of CDA patients’ red cells membrane abnormalities (55). Second, expression of *BCAM* gene is lower in individuals with the In(Lu) phenotype that is caused by a variety of KLF1 mutations that lead to its haploinsufficiency (18). Third, expression of *E2F2* is decreased in the presence of Nan-KLF1 mutation (24). Finally, the *Cdkn1B* gene (p27) is directly regulated by KLF1 during late stages of differentiation, where it is critical for enucleation of erythroid precursors (56) and this process is disturbed in the presence of CDA mutation (37,55). Furthermore, we expanded our analysis for *TFR2-* transferrin receptor 2, *SLC4A1*-anion transport protein (BAND3) and *IL17RB*-Interleukin-17 receptor B based on recently published article (55).

Analyses of RNA expression in the induced cells (Fig. 8*A*) show that *AQP1* and *EPB4.2* are not transactivated by CDA-KLF1 as efficiently as by WT-KLF1, resembling the data from CDA patients and in CDA-iPS-derived erythroblasts (Fig. 8*A*). On the other hand, *E2F2, BCAM, Cdkn1B, TFR2, SLC4A1* and *IL17RB* remain activated, and to an even greater level, by the CDA variant relative to WT. The expression of *IL17RB* is 8-fold higher with the presence in CDA-KLF1 than WT-KLF1 (Fig. 8*A*). As a potential explanation, inspection of the upstream promoters and introns of these genes reveals that the regulatory regions of *BCAM, Cdkn1B, E2F2, TFR2, SLC4A1* and *IL17RB* contain binding sites for both WT-KLF1 as well as CDA-KLF1, and thus are activated and transcribed (Fig. 8*C*). *AQP1* and *EPB4.2* contain sites only for WT-KLF1 that are not recognized by CDA-KLF1 (Fig. 8*B*), and thus unsuitable as an activation target. Interestingly, the *BCAM* result suggests a resolution of paradoxical patient observations, namely that *BCAM* expression is not affected in CDA patients while at the same time haploinsufficient levels of KLF1 are known to adversely affect *BCAM* expression. Our data suggest that the presence of variant DNA binding sites recognized by CDA-KLF1 protein compensates for any drop in expression of WT-KLF1 protein.

**Figure 8.**
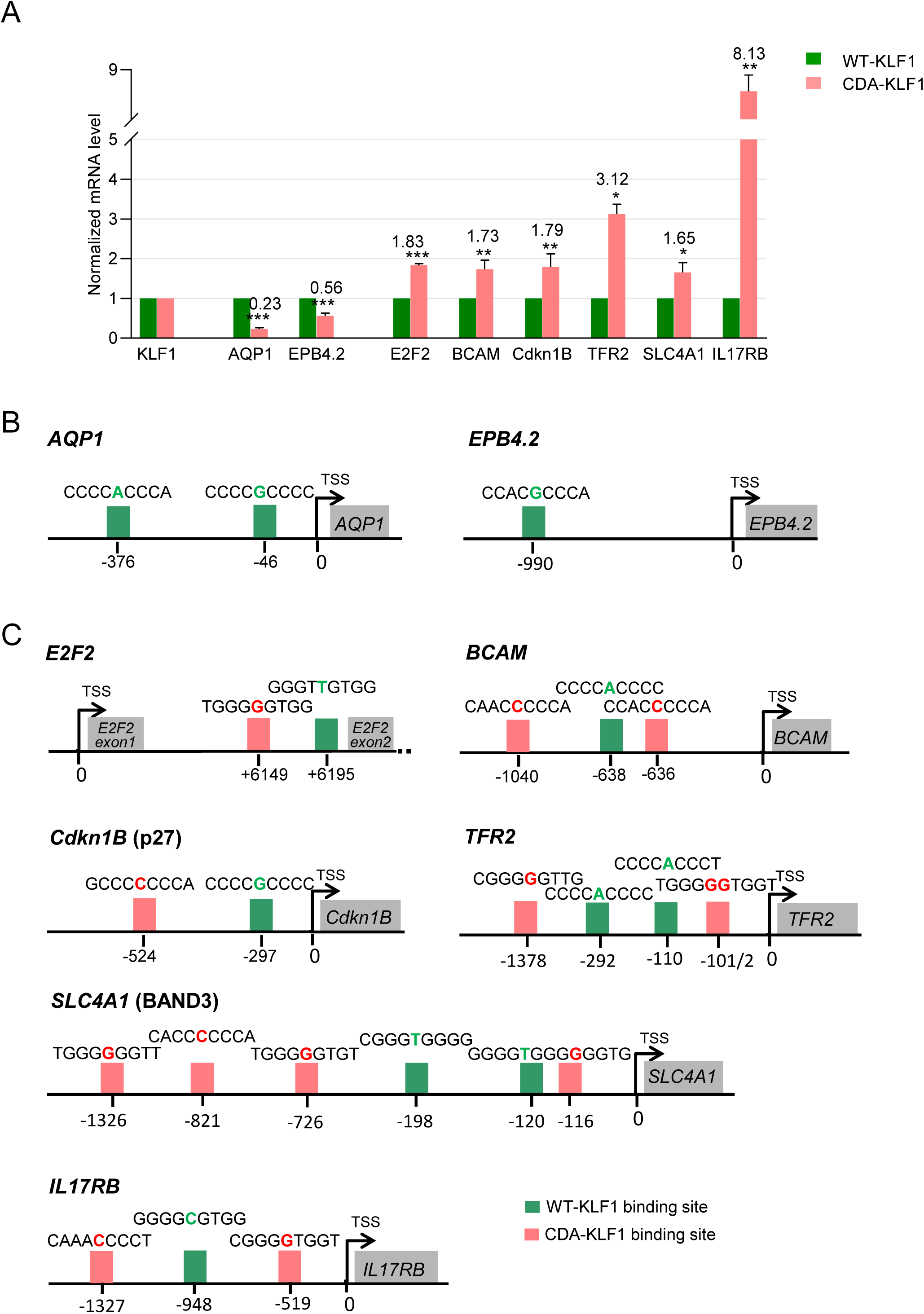
Quantitative analysis of the *in vivo* expression level of KLF1 target genes, carried out in stable K562 cell lines with induced expression of WT-KLF1 or CDA-KLF1. (*A*) RT-qPCR analyses of KLF1 target mRNA levels in stably transfected K562 cells with inducible expression of WT-KLF1 (green) and CDA-KLF1 (red). Data are normalized to GAPDH and to level of expression of WT-KLF1 or CDA-KLF1, respectively. Experiment was an average of triple repeats. (*B*) The scheme of human KLF1 target genes regulatory regions that contain only WT-KLF1 consensus binding sites marked in green, based on bioinformatics analysis. (*C*) The scheme of human KLF1 target genes regulatory regions that contain CDA-KLF1 consensus binding sites marked in red and WT-KLF1 binding sites marked in green, based on bioinformatics analysis. The location of binding sites based on distance from TSS. The individual DNA sequences of binding motifs are written above: *AQP1*-Aquaporin 1, *EPB4.2* - erythrocyte protein band 4.2, *E2F2*- E2F transcription factor 2, *BCAM*- basal cell adhesion molecule (Lutheran blood group), *Cdkn1B*- Cyclin Dependent Kinase Inhibitor 1B (p27*), TFR2*- transferrin receptor 2, *SLC4A1*- anion transport protein (BAND3) and *IL17RB* - Interleukin-17 receptor B. TSS-transcription start site. An average of at least three biological replicates of experiments (arithmetic mean ± SD) is shown *p≤0.05, ** p≤0.01, ***p≤ 0.001.

Collectively, these results show that CDA-KLF1 consensus binding site, as determined in the CASTing-Seq experiment, are uniquely recognized, bound, and transcriptionally functional endogenously in the cell.

## Discussion

Mutations of transcription factors that affect amino acids involved in direct interactions with DNA change their specificity. This leads to recognition and activation of an alternate set of genes and as a result may cause serious consequences for organisms. We have investigated such a mutation in KLF1, the essential transcription factor of erythropoiesis (reviewed in 21).

Using the *in vitro* CASTing method we identified a new set of sequences bound by CDA-KLF1, and based on them we defined the consensus binding site as 5’-NGG-G<u>G</u>T/G-T/GT/GT/G-3’. It differs from the consensus binding sites for WT-KLF1, 5’-NGG-G<u>C/T</u>G-T/GGG-3’ and for Nan-KLF1 5’-NGG-G<u>C/A</u>N-T/GGG-3’ as well. CDA-KLF1 binding site turns to be mutually exclusive compared to binding sites for WT-KLF1 and Nan-KLF1. It contains G in the middle (5^th^) position (underlined), whereas WT- KLF1 has C/T and Nan-KLF1 C/A. Our results are in agreement with the structural studies conducted by Carl Pabo, who analyzed the relationship between functionally important amino acids of zinc fingers and DNA sequence to which they bound (50,57,58). He found that if lysine is located in position +3 of the α-helix, within the ββα structure of the middle zinc finger, it has a high affinity for guanine in the middle (5^th^) position of the 9-nt binding site (50,57,58). Other research investigating YY1 and TFIIIA also described similar results, finding that lysine in zinc finger 2 of YY1 and in finger 1 of TFIIIA bound guanine (59-62). In addition, a detailed bioinformatics analysis of biochemical preferences between amino acids and bases in the DNA α-helix proves that lysine preferentially forms a strong bond with guanine (63), that entirely support our data.

Another interesting feature of our newly determined consensus binding site for CDA-KLF1 is high degeneracy of 3’end of the motif 5’-NGG-GG<u>T/G-T/GT/GT/G</u>-3’. On the G-rich strand, the last four positions of the motif could contain T or G nucleotides. Both bases, guanine and thymine can have alternate molecular structures based on different locations of a particular hydrogen atom. They can generate tautomeric keto or enol forms. Both guanine and thymine can switch easily from one tautomer to another. The change affects the three-dimensional shape of the molecule that may disturb the DNA: protein interface. The degeneracy of CDA-KLF1 consensus site generates a much wider combination of recognized sites as compared to WT-KLF1 or Nan-KLF1 that likely leads to a broader range of binding affinities for less specific targets. Occurrence of high degeneracy may be a solution for a too strong binding between transcription factor and target gene, which may translate to speed of the transcriptional reaction (16).

The results showing the CDA mutation located in ZnF2 affects binding of DNA sequence by ZnF1 were unexpected. We believe that the main reason for this effect are the high affinity of interactions between lysine (E339K) and “G” in the middle 5^th^ position of the 9-nt binding site. These interactions, along with the remaining interactions of ZnF2 and ZnF3 with G-rich positions 1^st^-4^th^ of the binding site may be sufficient to effectively bind CDA-KLF1 to DNA. Thus, the E to K mutation does not so much affect the specificity of ZnF1 interaction with DNA, which rather affects the binding strength/affinity of all three ZnFs, allowing for greater freedom and higher degeneration of the DNA sequence in interactions with ZnF1. The strong interactions in the complex would make the duration of binding too long, leading to low efficient function (64) as a transcription factor.

While this manuscript was prepared, the Perkins’ group published data on the consensus DNA-binding site of E325K KLF1 mutant using a different approach (65). Their consensus binding motif is in line with ours: 5’-NGG-G**<u>A</u>**<u>/</u>**<u>G</u>**G-T/GGG-3’. Binding affinity data show that the affinity of CDA-KLF1 for the site containing G in the 5^th^ position is approximately 5-fold higher than for sites with A (65). In addition, modeling studies based on the KLF4 zinc finger/DNA structure performed on a sequence with a central GGG triplet are consistent with our observations.

We tested the biochemical specificity between CDA-KLF1 in comparison with WT and Nan-KLF1 in complex formation with our defined DNA motif. We observed that the presence of G in the middle (5^th^) position of the binding motif entirely precludes complex formation with WT or Nan-KLF1. In the presence of CDA-KLF1 we detected a variation in the intensity of generated complexes suggesting that there are a range of affinities for the individual nucleotide sequences.

Both characteristics of the newly determined CDA-KLF1 consensus site (guanine in the middle position of the motif and the 3’end degeneracy) contribute to activation of neomorphic genes by CDA-KLF1 that are never activated by WT-KLF1. In the case of Nan-KLF1, the 6^th^ position of the consensus binding site 5’-NGG-GC/A<u>N</u>- T/GGG-3’ contains N that accepts any nucleotide. Such motif extends the range of recognizable binding sites relative to WT-KLF1, which only accepts G in this position. As a consequence, Nan-KLF1 activates neomorphic targets that are typically never expressed in the erythroid cells (27,28). The Nan-KLF1 mutant activates *Hamp* and *Irf7* genes (28). This ectopic gene expression contributes to the occurrence of chronic, lifelong anemia in the heterozygote Nan/+ mice (28). We can predict that a similar situation occurs in the case of CDA-KLF1 mutant. For example, patients suffering from CDA type IV present clinical manifestations outside of the erythroid system. Short stature and problems with urinary tract and gonad development are observed (26,29). Neomorphic gene activation by CDA-KLF1 may be one explanation for these characteristics.

Since we did not have access to CDA type IV patients, as this disease is very rare and only seven cases have been described (25,26,29-37), we prepared stable K562 cell lines with dox-inducible expression constructs of WT-KLF1 or CDA-KLF1. We investigated the level of transcription for known KLF1 target genes, which have both types of binding sites for the WT and CDA-KLF1 variants in their regulatory regions (promoters and/or introns). Such coexistence of both sites in close proximity may ameliorate the final outcome of gene activation for the CDA patients, as they are heterozygote for the mutation. CDA patients do not exhibit the In(Lu) phenotype (25,26,36,66), in which activation of the *Lu/BCAM* gene, the antigen of the Lutheran blood group, is sensitive to haploinsufficient levels of KLF1. Our analyses suggest that in CDA patients (that are genetically CDA/+), both WT and CDA KLF1 variants are able to bind and activate *Lu*/*BCAM* gene via independent promoter binding sites for both WT-KLF1 and CDA-KLF1. This clarifies previously published observations that were not entirely explained (16,67). This scenario may not be unique, as we found that the *E2F2* and *Cdkn1B* cell cycle regulator genes can be activated by both transcription factors.

It will be of interest to analyze the transcriptome of CDA type IV patients to verify that they indeed have ectopic expression of neomorphic, CDA-KLF1 activated genes. A recent study (55) shows that the genomic regions surrounding a number of the top ectopically expressed genes in a CDA patient (36) contain sites that, based on the present study, bind and are activated by CDA-KLF1 (5’-TGG-GGG-TGG-3’ (Fig. 1 and 7)). We tested some of them and the higher level of expression from CDA-KLF1 we obtained for IL17RB, that is critical for expression of IL8 (55). Since the genes activation by CDA-KLF1 were not confirmed by ChIP assay, we may not exclude the indirect effect of the presence of KLF1 mutant.

Activation of non-erythroid genes may help to explain the anomalies observed in CDA patients. It is also necessary to remember that zinc fingers of KLF1 are involved in interactions with proteins (9,68-73), such CDA mutation could disturb the interface between proteins and contribute to the pathological phenotype.

## Materials and Methods

### Plasmids and molecular cloning

For mammalian expression of Flag-tag KLF1 variants the pSG5-Flag-WT-KLF1 (68) plasmid was used and the mutations in amino acid position 339 of KLF1 were introduced by site directed mutagenesis (Agilent Technologies).

For bacterial expression of DBD of KLF1 variants the pMCSG7 vector was used. The Flag-tagged ZnF domains were subcloned from pSG5-Flag-WT-ZnF-KLF1 vector using BamHI restriction enzyme. The mutations in amino acid position 339 of KLF1 were introduced by site directed mutagenesis (Agilent Technologies), primers are available upon request. Luciferase reporter gene construction The pGL3 Basic vector (Promega) was used as a backbone. The 63 nucleotides long HSV thymidine kinase minimal promoter (TK mini) synthesized by SIGMA-ALDRICH, was added upstream of luciferase gene. To such construct, pGL3-TKmini, six repeats of CASTing selected binding sites synthesized by SIGMA-ALDRICH were introduced upstream of the TKmini promoter. Primers used to insert six repeats of indicated DNA binding sites are available upon request.

### Cell lines, transfection, nuclear extracts isolation and western blot analysis

K562 cells were cultured in RPMI medium 1640 supplemented with 10% fetal bovine serum (Corning), 0.5% Penicillin-Streptomycin (BioShop). COS-7 cell lines were cultured in Dulbecco’s modified Eagle’s medium supplemented with 10% fetal bovine serum, 0.5% Penicillin-Streptomycin. All cells were cultured in a humidified incubator at 37°C in a 5% CO2 and 95% air atmosphere. To prepare protein extracts for EMSA, COS-7 cells growing on 10cm diameter dishes were transfected with 10μg of plasmids: pSG5-WT-KLF1, Nan or CDA using Lipofectamine LTX according to the manufacturer’s protocol. After 48 h of incubation, crude extracts were prepared as described previously (74). For western blot analysis of Flag-tagged KLF1 variants, PVDF membranes were probed with anti-Flag-tag Monoclonal M2-Peroxidase (HRP) antibody (SIGMA) or anti-KLF1 7B2 antibody as was described (13).

### Recombinant proteins expression and purification

KLF1 Flag-tag zinc finger domains: CDA-ZnF-KLF1 and WT-ZnF-KLF1 were cloned into pMCSG7 to generate an in-frame fusion to 6x-His-Tag and TEV cleavage sites. The CDA mutation (E to K) was introduced to mouse ZnF domain in position 339 aa by site directed mutagenesis, according to manufacture protocol (Agilent Technologies). Primers for mutagenesis are listed in Table 1.

**Table 1.**
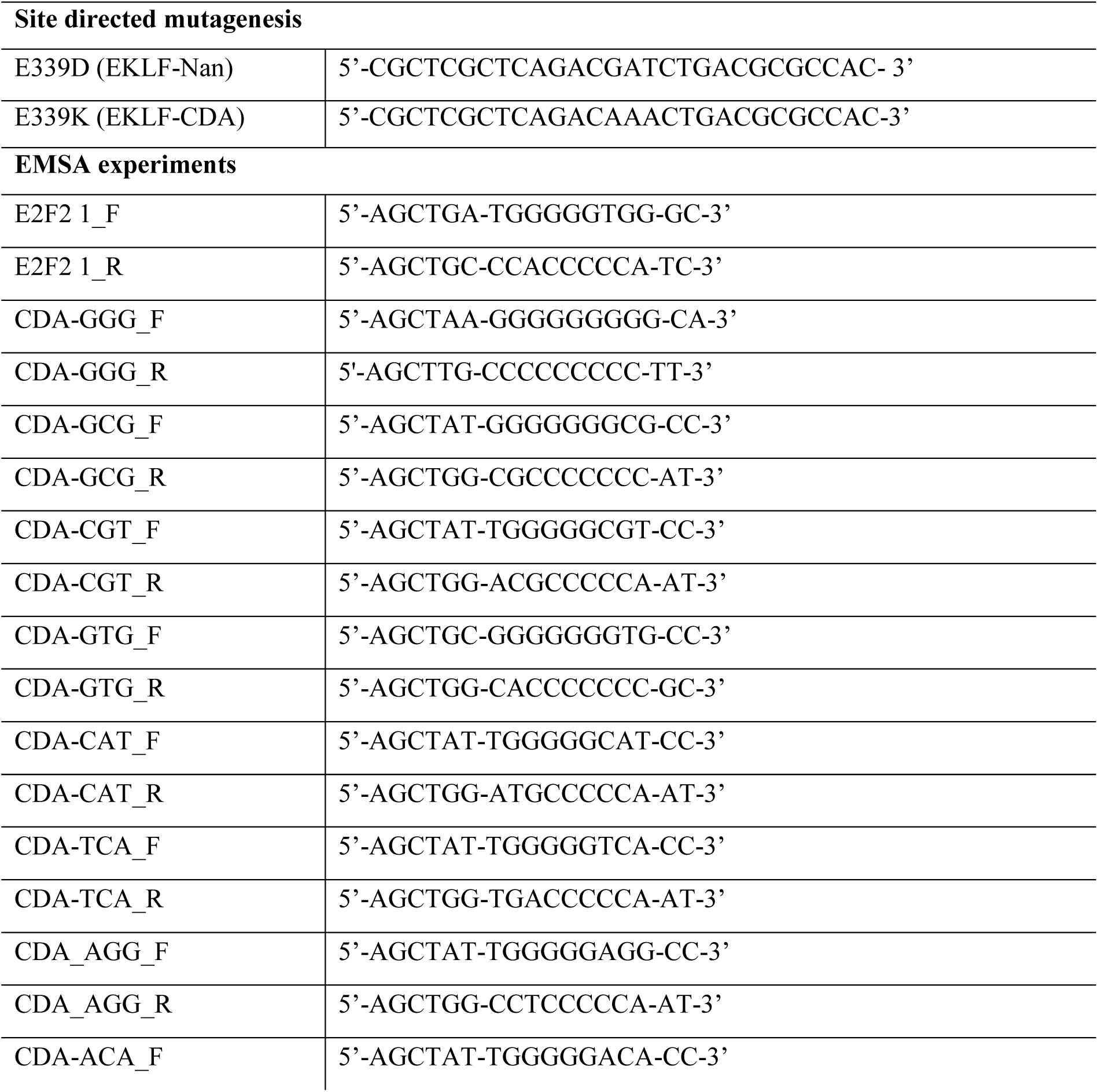
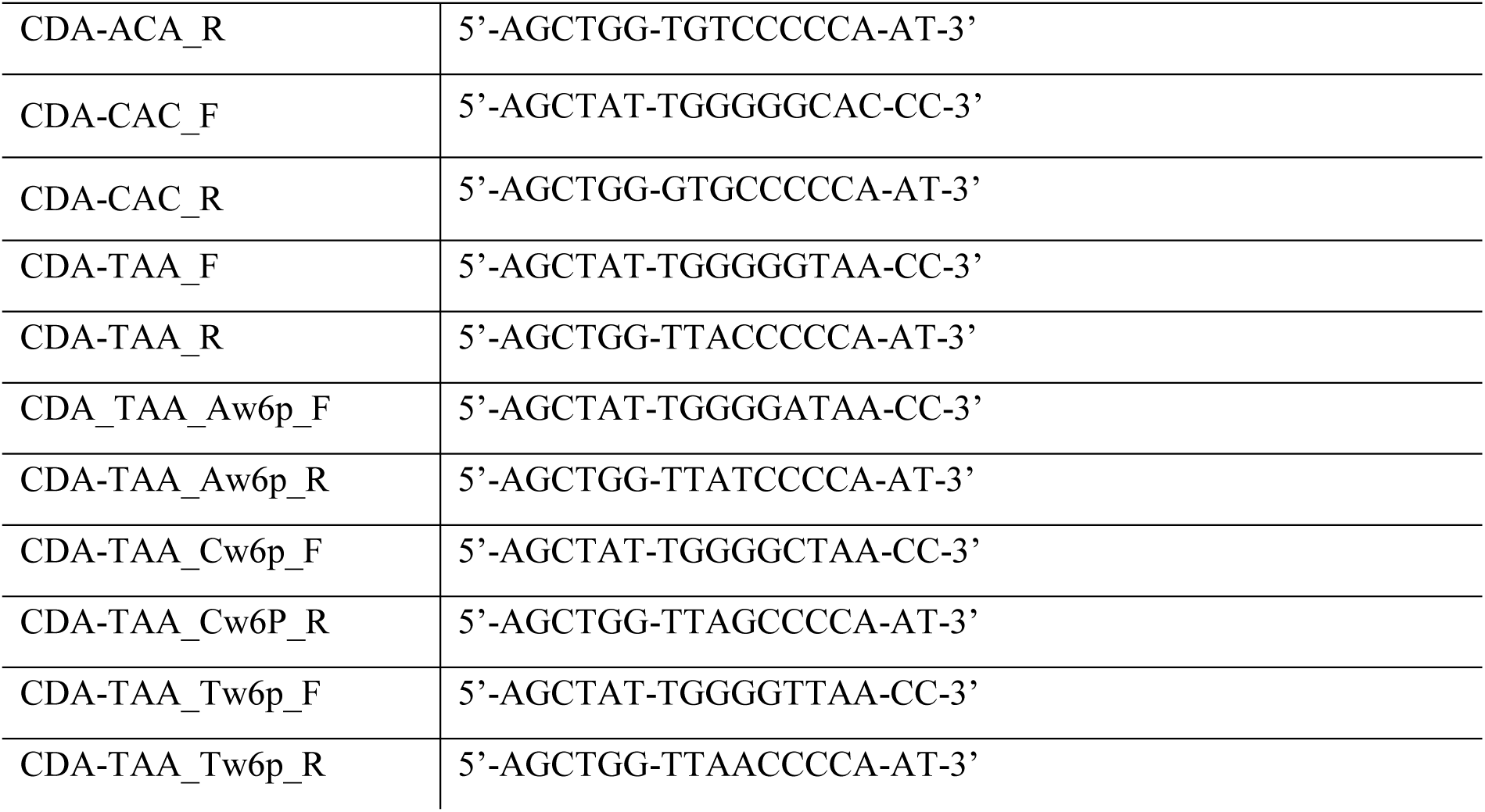
Primer sequences used in this study.

To purified DBD of KLF1 variants the pMCSG7 plasmid containing Flag-CDA-ZnF-KLF1 and FlagWT-ZnF-KLF1 were used. BL21-CodonPlus(DE3)-RIL E. coli were transformed with these plasmids and induced with 0.3 mM isopropyl β-D-1-thiogalactopyranoside (IPTG). Bacteria were cultured in the presence of 100µM ZnCl2 to assist in protein folding for 17h at 18°C before collection. Cell pellets were resuspended in cold lysis buffer (50mM Tris-HCL pH 8.0, 1.5M NaCl, 10% glycerol and 0.5mM TCEP) with addition of sarcosyl to final 2%. Cell lysates were sonicated for 4 minutes (2 sec ON, 20 sec OFF) and centrifuged at 30500g for 30 min at 4°C. Supernatants were diluted ten times in lysis S-2 buffer and loaded on column with Zn-resin equilibrated with lysis buffer (75). Column was washed with lysis buffer with increasing salt concentration (150mM-1.5M). Then the salt concentration was decreased to 150mM allowing for efficient, on column, TEV protease cleavage for 2h at room temperature. 120µg TEV protease for every 1mg of protein was used. Recombinant zinc finger domains were eluted from column with elution buffer (50mM Tris-HCl, 150mM NaCl, 10% glycerol, 1M NH4Cl). Fractions containing recombinant DBD were combined and purified on Superdex HiLoad 200 column equilibrated with buffer (50mM Hepes pH7.9, 100mM KCl, 10% glycerol, 10µM ZnCl2, 0.5mM TCEP) on an automated FPLC system (AKTApurifier). Chromatography was carried out at 4°C.

### CASTing for the CDA-KLF1 consensus DNA-binding site

CASTing was performed as previously described (41,44) with some modifications. In first CASTing experiment we used the library with twelve random nucleotides:

5’-CAAGCTTACTGCAGATGCNNNNNNNNNNNNCGTAGGATCCATCTAGAGT-3’ flanked by constant sequences needed for amplification; in second attempt we used the library consisted of 8 random nucleotides with fixed G in the middle position: 5’-GCTCAAGCTTACTGCAGATATNNNNGNNNNTTTAGGATCCATCTAGAGTCCGA- 3’(SIGMA-ALDRICH). 15µg of the library were made double-stranded with 5µg 3’ primer (CAST1_R- ACTCTAGATGGATCCTACG or CAST2_R-TCGGACTCTAGATGGATCCTA) in a reaction that contained 200 mM each of the four deoxynucleoside triphosphates and 20 U DreamTaq DNA polymerase in 1x DreamTaq Buffer (ThermoScientific) for one cycle of 98°C for 3 min, 94°C for 1 min, 47°C for 2 min and 72°C for 30 min. CASTing procedure: 12µl of ANTI-FLAG® M2 Magnetic Beads (SIGMA-ALDRICH) were washed twice in 40µl of ESB buffer (20mM HEPES [pH 7.9], 40mM KCl, 6mM MgCl2, 1mM dithiothreitol, 1mM phenylmethylsulfonyl fluoride, 0.1% NP-40, 10% glycerol), then 800ng of purified ZnF-KLF1 (WT or CDA) protein in 12µl of Binding Buffer (10mM Tris–HCl, pH 7.5, 150mM NaCl, 20 mM KCl, 10mM MgCl2 and 5mM ZnCl2) was added and incubated for 1 hour in 4°C. Immunoprecipitates were washed three times in EBC buffer (50mM Tris-HCL pH 8.0, 120mM NaCl, 0.5% NP-40, 5mM NaF, 1mM Na3VO4, 10µg/ml phenylmethylsulfonyl fluoride, 10µg/ml leupeptin) and twice in ESB buffer. Next, beads with attached ZnF domains were suspended in 25µl ESB buffer with 150ng sonicated salmon sperm DNA (sssDNA, Fermentas) and 150ng of the double stranded library was added. The mixture was incubated for 30 minutes at room temperature on a low-speed shaking platform. The beads/protein/ds DNA oligo complexes were washed three times in ESB buffer and resulted complexes served as a template for amplification in 50µl of PCR reaction mixture containing 200ng of CAST_F and CAST_R primers, 0.5mCi of [α-32P]dCTP, 20mM dCTP, 50mM each dATP, dTTP, dGTP, 10µg of bovine serum albumin, 1x Dream Taq Buffer, and 1µl of Dream Taq polymerase (Thermo Scientific). Selected ds DNA oligos were amplified for 15 cycles of 1 min at 94°C, 1 min at 62°C, 1 min at 72°C and 7 min for final extension. Amplified dsDNA oligos were purified on a Sephadex G-50 spin column (GE Health Care) and resolved on 8% nondenaturing polyacrylamide gels. Radiolabeled oligonucleotides were excised from gels, eluted overnight, and used in subsequent CASTing cycles. After five rounds of selection, adaptors and barcodes for NGS sequencing were added by fusion PCR (primers available upon request). The quantity and quality of the pool of selected dsDNA oligos were determined using Qubit fluorometric assay and Agilent BioAnalyzer High-Sensitivity DNA kit (Agilent Technologies). Emulsion PCR and Ion Sphere Particles (ISP) enrichment were done using the Ion PGM Hi-Q Kit for 200bp (Life Technologies) according to the manufacturer’s instructions. Next generation sequencing was carried out on Ion Torrent Personal Genome Machine sequencer (Life Technologies) using Ion 314 Chip (Life Technologies,) and Ion PGM Hi-Q Sequencing Kit (Life Technologies) according to the manufacturer’s instructions. The obtained ∼ 30000 sequences were analyzed by the MEME software to perform de novo motif discovery (76).

### Electro-mobility shift assay (EMSA)

The gel retardation assay was performed as previously described (24) with some modifications, briefly: double-stranded synthetic oligonucleotides containing specific KLF1 binding sites found by CASTing were radiolabeled by [α-32P]-CTP using Klenow polymerase and used as a probes or competitors for gel retardation analysis. Binding reactions were performed in the presence of ∼ 10μg (5µg/µl) of the whole-cell extracts from COS-7 cells transfected with WT, CDA or Nan-KLF1 variants or purified recombinant ZnF domains of WT or CDA-KLF1. The mix was incubated for 20 minutes in Binding Buffer (25mM HEPES, pH 7.5, 32mM KCI, 50mM NaCl, 2µM ZnCl2, 0.7mM β-mercaptoethanol, 8% glycerol) on ice (whole extract) or RT(recombinant protein). Unspecific interactions were quenched with 100ng/µl (whole extract) or 40ng/µl (recombinant protein) of sssDNA (Fermentas) and resolved in a non-denaturing 8% polyacrylamide gel in 0.5X TBE buffer at 4°C. Anti-KLF1 antibodies 4B9 were used to identify the KLF1 band position (7). For quantitative measurement of dissociation constant the following concentration of recombinant WT or CDA-ZnF-KLF1 domain were used: 0, 100, 150, 500, 750, 980, 1230, 1470, 1720nM. Curves were fit using Hill Slope in GraphPad. Averaged Kd and its standard deviation were calculated based on at least two EMSA gel shifts. Oligonucleotide sequences used in EMSA are listed in Table 1.

### Reporter Gene Analysis-Dual Luciferase Assays

K562 cells seeded in 24-well plates were transfected with suitable reporter genes and increasing amount of plasmid carrying WT, Nan or CDA-KLF1 using Lipofectamine LTX. After 36h of incubation cells were lysed and assayed for luciferase activities with a dual luciferase system (Promega). Plasmid pRLTK (Renilla) Promega was included as a normalization control for transfection efficiency. Luminescence was quantified with a luminometer (GloMax 96-microplate, Promega). The results are the average of at least three experiments performed in triplicates.

### Generation of stable K562 cell lines

Plasmids pSG5 containing mouse Flag-WT/CDA-KLF1 serve as a sources of full length of KLF1 variant genes which were amplified by PCR and introduced into the pSAM vector (kind gift from Rita Perlingeiro, PhD University of Minnesota). For lentiviral production, HEK-293T cells were cotransfected with gene-of-interest (GOI)-containing vectors with doxycycline dependent promoter and components of 2nd generation packaging vectors: pPAX2 packaging vector and pMD2G envelope vector using linear PEI 25000 (SIGMA-ALDRICH). 48 h post-transfection, the lentiviruses were collected, filtered and added to the K562 cells. After 72 h the expression of KLF variants were induced by 1µg/ml doxycycline. To select KLF1 variants containing cells sorting for fluorescent-positive cells using a FACSAria III cell sorter (BD Biosciences, La Jolla, CA, USA) was performed. The primers used to insert KLF1 variants into pSAM vectors are available upon request.

### RNA isolation and Real-time PCR analyses

Total RNA was extracted from stable K562 cell lines with induced expression of WT or CDA-KLF1 (1.5µg/ml doxycycline for 18h) using 3-zone® Reagent (Novazym) according to the manufacturer’s instructions. Extracted RNA was treated with DNase I (Thermofisher Scientific) and subjected to cDNA synthesis using a M-MLV Reverse Transcriptase kit from Verte according to the manufacturer’s instructions (Novazym). Gene expression was quantified using Real-time (RT-qPCR) with SsoAdvanced™ Universal SYBR® Green Supermix on a CFX Connect™ Real-Time PCR Detection System (BioRAD). The primers used for RT-qPCR are available upon request.

## Acknowledgements

We are grateful for help in: protein purification Dr. Barbara Imiolczyk (Inst. of Bioorganic Chemistry, PAS), bioinformatics analyses Alexey Bryzgalov, Prof. Izabela Makalowska from Adam Mickiewicz University (AMU), generation of stable cell lines Anna Witucka (AMU), Dr. Jacek Stepniewki, Prof. Alicja Jozkowicz and Prof. Jozef Dulak (Jagiellonian Uni., Krakow). Molecular Biology Techniques Laboratory (AMU) for NGS. This work was supported by the Polish National Science Center 2013/09/B/NZ1/01879 to MS and by the National Institutes of Health DK046865 to JJB.

## Abbreviations

AQP1: Aquaporin 1
CASTing: Cyclic Amplification and Selection of Targets
CDA: Congenital Dyserythropoietic Anemia
Cdkn1B: Cyclin Dependent Kinase Inhibitor 1B (p27)
COS-7: African green monkey fibroblast
E2F2: E2F transcription factor 2
EMSA: Electrophoretic mobility shift assay
EPB4.2: erythrocyte protein band 4.2
HPFH: Hereditary Persistence of Fetal Hemoglobin
IL17RB: Interleukin-17 receptor B
IPTG: Isopropyl β-D-1-thiogalactopyranoside
K562: erythroleukemia type cells
KLF: Krüppel-like Factor
Lu/BCAM: basal cell adhesion molecule (Lutheran blood group)
Nan: Neonatal anemia
P/CAF: P300/CBP-associated factor
CBP: CREB1-binding protein
SLC4A1: anion transport protein (BAND3)
SWI/SNF: SWItch/Sucrose Non-Fermentable
TFIIIA: transcription factor IIIA
TFR2: transferrin receptor 2
TSS: transcription start site
YY1: Yin Yang 1 transcriptional repressor protein
ZnF: zinc finger

## References

1. Miller, I. J., and Bieker, J. J. (1993) A novel, erythroid cell-specific murine transcription factor that binds to the CACCC element and is related to the Kruppel family of nuclear proteins. Mol Cell Biol 13, 2776–2786

2. Jenkins, N. A., Gilbert, D. J., Copeland, N. G., Gruzglin, E., and Bieker, J. J. (1998) Erythroid Kruppel-like transcription factor (Eklf) maps to a region of mouse chromosome 8 syntenic with human chromosome 19. Mamm Genome 9, 174–176

3. Pilon, A. M., Ajay, S. S., Kumar, S. A., Steiner, L. A., Cherukuri, P. F., Wincovitch, S., Anderson, S. M., Comparative Sequencing Program, N., Mullikin, J. C., Gallagher, P. G., Hardison, R. C., Margulies, E. H., and Bodine, D. M. (2011) Genome-wide ChIP-Seq reveals a dramatic shift in the binding of the transcription factor erythroid Kruppel-like factor during erythrocyte differentiation. Blood

4. Tallack, M. R., Whitington, T., Shan Yuen, W., Wainwright, E. N., Keys, J. R., Gardiner, B. B., Nourbakhsh, E., Cloonan, N., Grimmond, S. M., Bailey, T. L., and Perkins, A. C. (2010) A global role for KLF1 in erythropoiesis revealed by ChIP-seq in primary erythroid cells Genome Research 20 1052–1063

5. Perkins, A. C., Sharpe, A. H., and Orkin, S. H. (1995) Lethal beta-thalassaemia in mice lacking the erythroid CACCC-transcription factor EKLF. Nature 375, 318–322

6. Nuez, B., Michalovich, D., Bygrave, A., Ploemacher, R., and Grosveld, F. (1995) Defective haematopoiesis in fetal liver resulting from inactivation of the EKLF gene. Nature 375, 316–318

7. Zhang, W., Kadam, S., Emerson, B. M., and Bieker, J. J. (2001) Site-specific acetylation by p300 or CREB binding protein regulates erythroid Kruppel-like factor transcriptional activity via its interaction with the SWI-SNF complex. Mol Cell Biol 21, 2413–2422

8. Armstrong, J. A., Bieker, J. J., and Emerson, B. M. (1998) A SWI/SNF-related chromatin remodeling complex, E-RC1, is required for tissue-specific transcriptional regulation by EKLF in vitro. Cell 95, 93–104

9. Kadam, S., McAlpine, G. S., Phelan, M. L., Kingston, R. E., Jones, K. A., and Emerson, B. M. (2000) Functional selectivity of recombinant mammalian SWI/SNF subunits. Genes Dev 14, 2441–2451

10. Yien, Y. Y., and Bieker, J. J. (2013) EKLF/KLF1, a tissue-restricted integrator of transcriptional control, chromatin remodeling, and lineage determination. Mol Cell Biol 33, 4–13

11. Drissen, R., Palstra, R. J., Gillemans, N., Splinter, E., Grosveld, F., Philipsen, S., and de Laat, W. (2004) The active spatial organization of the beta-globin locus requires the transcription factor EKLF. Genes Dev 18, 2485–2490

12. Ouyang, L., Chen, X., and Bieker, J. J. (1998) Regulation of erythroid Kruppel-like factor (EKLF) transcriptional activity by phosphorylation of a protein kinase casein kinase II site within its interaction domain. J Biol Chem 273, 23019–23025

13. Siatecka, M., Xue, L., and Bieker, J. J. (2007) Sumoylation of EKLF Promotes Transcriptional Repression and Is Involved in Inhibition of Megakaryopoiesis Mol. Cell. Biol. 27, 8547–8560

14. Zhang, W., and Bieker, J. J. (1998) Acetylation and modulation of erythroid Kruppel-like factor (EKLF) activity by interaction with histone acetyltransferases. Proc Natl Acad Sci U S A 95, 9855–9860

15. Quadrini, K. J., and Bieker, J. J. (2006) EKLF/KLF1 is ubiquitinated in vivo and its stability is regulated by activation domain sequences through the 26S proteasome. FEBS Lett 580, 2285–2293

16. Singleton, B. K., Lau, W., Fairweather, V. S., Burton, N. M., Wilson, M. C., Parsons, S. F., Richardson, B. M., Trakarnsanga, K., Brady, R. L., Anstee, D. J., and Frayne, J. (2011) Mutations in the second zinc finger of human EKLF reduce promoter affinity but give rise to benign and disease phenotypes. Blood 118, 3137–3145

17. Perkins, A., Xu, X., Higgs, D. R., Patrinos, G. P., Arnaud, L., Bieker, J. J., Philipsen, S., and Workgroup, K. L. F. C. (2016) Kruppeling erythropoiesis: an unexpected broad spectrum of human red blood cell disorders due to KLF1 variants. Blood 127, 1856–1862

18. Singleton, B. K., Burton, N. M., Green, C., Brady, R. L., and Anstee, D. J. (2008) Mutations in EKLF/KLF1 form the molecular basis of the rare blood group In(Lu) phenotype. Blood 112, 2081–2088

19. Borg, J., Patrinos, G. P., Felice, A. E., and Philipsen, S. (2011) Erythroid phenotypes associated with KLF1 mutations. Haematologica 96, 635–638

20. Liu, D., Zhang, X., Yu, L., Cai, R., Ma, X., Zheng, C., Zhou, Y., Liu, Q., Wei, X., Lin, L., Yan, T., Huang, J., Mohandas, N., An, X., and Xu, X. (2014) KLF1 mutations are relatively more common in a thalassemia endemic region and ameliorate the severity of beta-thalassemia. Blood 124, 803–811

21. Siatecka, M., and Bieker, J. J. (2011) The multifunctional role of EKLF/KLF1 during erythropoiesis. Blood 118, 2044–2054

22. Heruth, D. P., Hawkins, T., Logsdon, D. P., Gibson, M. I., Sokolovsky, I. V., Nsumu, N. N., Major, S. L., Fegley, B., Woods, G. M., Lewing, K. B., Neville, K. A., Cornetta, K., Peterson, K. R., and White, R. A. (2010) Mutation in erythroid specific transcription factor KLF1 causes Hereditary Spherocytosis in the Nan hemolytic anemia mouse model. Genomics 96, 303–307

23. White, R. A., Sokolovsky, I. V., Britt, M. I., Nsumu, N. N., Logsdon, D. P., McNulty, S. G., Wilmes, L. A., Brewer, B. P., Wirtz, E., Joyce, H. R., Fegley, B., Smith, A., and Heruth, D. P. (2009) Hematologic characterization and chromosomal localization of the novel dominantly inherited mouse hemolytic anemia, neonatal anemia (Nan). Blood Cells Mol Dis 43, 141–148

24. Siatecka, M., Sahr, K. E., Andersen, S. G., Mezei, M., Bieker, J. J., and Peters, L. L. (2010) Severe anemia in the Nan mutant mouse caused by sequence-selective disruption of erythroid Kruppel-like factor. Proc Natl Acad Sci U S A 107, 15151–15156

25. Singleton, B. K., Fairweather, V. S., Lau, W., Parsons, S. F., Burton, N. M., Frayne, J., Brady, R. L., and Anstee, D. J. (2009) A Novel EKLF Mutation in a Patient with Dyserythropoietic Anemia: The First Association of EKLF with Disease in Man. Blood 114, 162–162

26. Arnaud, L., Saison, C., Helias, V., Lucien, N., Steschenko, D., Giarratana, M.-C., Prehu, C., Foliguet, B., Montout, L., de Brevern, A. G., Francina, A., Ripoche, P., Fenneteau, O., Da Costa, L., Peyrard, T., Coghlan, G., Illum, N., Birgens, H., Tamary, H., Iolascon, A., Delaunay, J., Tchernia, G., and Cartron, J.-P. (2010) A Dominant Mutation in the Gene Encoding the Erythroid Transcription Factor KLF1 Causes a Congenital Dyserythropoietic Anemia. The American Journal of Human Genetics 87, 721–727

27. Gillinder, K. R., Ilsley, M. D., Nebor, D., Sachidanandam, R., Lajoie, M., Magor, G. W., Tallack, M. R., Bailey, T., Landsberg, M. J., Mackay, J. P., Parker, M. W., Miles, L. A., Graber, J. H., Peters, L. L., Bieker, J. J., and Perkins, A. C. (2017) Promiscuous DNA-binding of a mutant zinc finger protein corrupts the transcriptome and diminishes cell viability. Nucleic Acids Res 45, 1130–1143

28. Planutis, A., Xue, L., Trainor, C. D., Dangeti, M., Gillinder, K., Siatecka, M., Nebor, D., Peters, L. L., Perkins, A. C., and Bieker, J. J. (2017) Neomorphic effects of the neonatal anemia (Nan-Eklf) mutation contribute to deficits throughout development. Development 144, 430–440

29. Ravindranath, Y., Johnson, R. M., Goyette, G., Buck, S., Gadgeel, M., and Gallagher, P. G. (2018) KLF1 E325K-associated Congenital Dyserythropoietic Anemia Type IV: Insights Into the Variable Clinical Severity. J Pediatr Hematol Oncol 40, e405–e409

30. Ortolano, R., Forouhar, M., Warwick, A., and Harper, D. (2018) A Case of Congenital Dyserythropoeitic Anemia Type IV Caused by E325K Mutation in Erythroid Transcription Factor KLF1. J Pediatr Hematol Oncol 40, e389–e391

31. de-la-Iglesia-Inigo, S., Moreno-Carralero, M. I., Lemes-Castellano, A., Molero-Labarta, T., Mendez, M., and Moran-Jimenez, M. J. (2017) A case of congenital dyserythropoietic anemia type IV. Clin Case Rep 5, 248–252

32. Tang, W., Cai, S. P., Eng, B., Poon, M. C., Waye, J. S., Illum, N., and Chui, D. H. (1993) Expression of embryonic zeta-globin and epsilon-globin chains in a 10-year-old girl with congenital anemia. Blood 81, 1636–1640

33. Agre, P., Asimos, A., Casella, J. F., and McMillan, C. (1986) Inheritance pattern and clinical response to splenectomy as a reflection of erythrocyte spectrin deficiency in hereditary spherocytosis. N Engl J Med 315, 1579–1583

34. Wickramasinghe, S. N., Illum, N., and Wimberley, P. D. (1991) Congenital dyserythropoietic anaemia with novel intra-erythroblastic and intra-erythrocytic inclusions. Br J Haematol 79, 322–330

35. Parsons, S. F., Jones, J., Anstee, D. J., Judson, P. A., Gardner, B., Wiener, E., Poole, J., Illum, N., and Wickramasinghe, S. N. (1994) A novel form of congenital dyserythropoietic anemia associated with deficiency of erythroid CD44 and a unique blood group phenotype [In(a-b-), Co(a-b-)]. Blood 83, 860–868

36. Jaffray, J. A., Mitchell, W. B., Gnanapragasam, M. N., Seshan, S. V., Guo, X., Westhoff, C. M., Bieker, J. J., and Manwani, D. (2013) Erythroid transcription factor EKLF/KLF1 mutation causing congenital dyserythropoietic anemia type IV in a patient of Taiwanese origin: review of all reported cases and development of a clinical diagnostic paradigm. Blood Cells Mol Dis 51, 71–75

37. Kohara, H., Utsugisawa, T., Sakamoto, C., Hirose, L., Ogawa, Y., Ogura, H., Sugawara, A., Liao, J., Aoki, T., Iwasaki, T., Asai, T., Doisaki, S., Okuno, Y., Muramatsu, H., Abe, T., Kurita, R., Miyamoto, S., Sakuma, T., Shiba, M., Yamamoto, T., Ohga, S., Yoshida, K., Ogawa, S., Ito, E., Kojima, S., Kanno, H., and Tani, K. (2019) KLF1 mutation E325K induces cell cycle arrest in erythroid cells differentiated from congenital dyserythropoietic anemia patient-specific induced pluripotent stem cells. Exp Hematol 73, 25–37 e28

38. Renella, R., and Wood, W. G. (2009) The congenital dyserythropoietic anemias. Hematol Oncol Clin North Am 23, 283–306

39. Iolascon, A., Russo, R., and Delaunay, J. (2011) Congenital dyserythropoietic anemias. Curr Opin Hematol 18, 146–151

40. Heimpel, H., Wendt, F., Klemm, D., Schubothe, H., and Heilmeyer, L. (1968) [Congenital dyserythropoietic anemia]. Arch Klin Med 215, 174–194

41. Tao, Y., Kassatly, R. F., Cress, W. D., and Horowitz, J. M. (1997) Subunit composition determines E2F DNA-binding site specificity. Mol Cell Biol 17, 6994–7007

42. Corbi, N., Perez, M., Maione, R., and Passananti, C. (1997) Synthesis of a new zinc finger peptide; comparison of its ‘code’ deduced and ‘CASTing’ derived binding sites. FEBS Lett 417, 71–74

43. Thiesen, H. J., and Bach, C. (1990) Target Detection Assay (TDA): a versatile procedure to determine DNA binding sites as demonstrated on SP1 protein. Nucleic Acids Res 18, 3203–3209

44. Shields, J. M., and Yang, V. W. (1998) Identification of the DNA sequence that interacts with the gut-enriched Kruppel-like factor. Nucleic Acids Res 26, 796–802

45. Gu, G., Wang, T., Yang, Y., Xu, X., and Wang, J. (2013) An improved SELEX-Seq strategy for characterizing DNA-binding specificity of transcription factor: NF-kappaB as an example. PLoS One 8, e76109

46. Klevit, R. E. (1991) Recognition of DNA by Cys2, His2 zinc fingers. Science 253, 1367, 1393

47. Feng, W. C., Southwood, C. M., and Bieker, J. J. (1994) Analyses of beta-thalassemia mutant DNA interactions with erythroid Kruppel-like factor (EKLF), an erythroid cellspecific transcription factor. J Biol Chem 269, 1493–1500

48. Pabo, C. O., and Sauer, R. T. (1992) Transcription factors: structural families and principles of DNA recognition. Annu Rev Biochem 61, 1053–1095

49. Tallack, M. R., Keys, J. R., Humbert, P. O., and Perkins, A. C. (2009) EKLF/KLF1 Controls Cell Cycle Entry via Direct Regulation of E2f2 Journal of Biological Chemistry 284 20966–20974

50. Wolfe, S. A., Nekludova, L., and Pabo, C. O. (2000) DNA recognition by Cys2His2 zinc finger proteins. Annu Rev Biophys Biomol Struct 29, 183–212

51. Siatecka, M., Lohmann, F., Bao, S., and Bieker, J. J. (2010) EKLF Directly Activates the p21WAF1/CIP1 Gene by Proximal Promoter and Novel Intronic Regulatory Regions during Erythroid Differentiation. Mol. Cell. Biol. 30, 2811–2822

52. Donze, D., Townes, T. M., and Bieker, J. J. (1995) Role of erythroid Kruppel-like factor in human gamma-to beta-globin gene switching. J Biol Chem 270, 1955–1959

53. Orkin, S. H., Antonarakis, S. E., and Kazazian, H. H., Jr. (1984) Base substitution at position −88 in a beta-thalassemic globin gene. Further evidence for the role of distal promoter element ACACCC. J Biol Chem 259, 8679–8681

54. Agre, P., Smith, B. L., Baumgarten, R., Preston, G. M., Pressman, E., Wilson, P., Illum, N., Anstee, D. J., Lande, M. B., and Zeidel, M. L. (1994) Human red cell Aquaporin CHIP. II. Expression during normal fetal development and in a novel form of congenital dyserythropoietic anemia. J Clin Invest 94, 1050–1058

55. Varricchio, L., Planutis, A., Manwani, D., Jaffray, J., Mitchell, W. B., Migliaccio, A. R., and Bieker, J. J. (2019) Genetic disarray follows mutant KLF1-E325K expression in a congenital dyserythropoietic anemia patient. Haematologica

56. Gnanapragasam, M. N., McGrath, K. E., Catherman, S., Xue, L., Palis, J., and Bieker, J. J. (2016) EKLF/KLF1-regulated cell cycle exit is essential for erythroblast enucleation. Blood 128, 1631–1641

57. Pavletich, N. P., and Pabo, C. O. (1991) Zinc finger-DNA recognition: crystal structure of a Zif268-DNA complex at 2.1 A. Science 252, 809–817

58. Wolfe, S. A., Greisman, H. A., Ramm, E. I., and Pabo, C. O. (1999) Analysis of zinc fingers optimized via phage display: evaluating the utility of a recognition code. J Mol Biol 285, 1917–1934

59. Houbaviy, H. B., Usheva, A., Shenk, T., and Burley, S. K. (1996) Cocrystal structure of YY1 bound to the adeno-associated virus P5 initiator. Proc Natl Acad Sci U S A 93, 13577–13582

60. Wuttke, D. S., Foster, M. P., Case, D. A., Gottesfeld, J. M., and Wright, P. E. (1997) Solution structure of the first three zinc fingers of TFIIIA bound to the cognate DNA sequence: determinants of affinity and sequence specificity. J Mol Biol 273, 183–206

61. Nolte, R. T., Conlin, R. M., Harrison, S. C., and Brown, R. S. (1998) Differing roles for zinc fingers in DNA recognition: structure of a six-finger transcription factor IIIA complex. Proc Natl Acad Sci U S A 95, 2938–2943

62. Segal, D. J., Dreier, B., Beerli, R. R., and Barbas, C. F., 3rd. (1999) Toward controlling gene expression at will: selection and design of zinc finger domains recognizing each of the 5’-GNN-3’ DNA target sequences. Proc Natl Acad Sci U S A 96, 2758–2763

63. Luscombe, N. M., Laskowski, R. A., and Thornton, J. M. (2001) Amino acid-base interactions: a three-dimensional analysis of protein-DNA interactions at an atomic level. Nucleic Acids Res 29, 2860–2874

64. Chu, X., and Wang, J. (2014) Specificity and Affinity Quantification of Flexible Recognition from Underlying Energy Landscape Topography. PLOS Computational Biology 10, e1003782

65. Ilsley, M. D., Huang, S., Magor, G. W., Landsberg, M. J., Gillinder, K. R., and Perkins, A. C. (2019) Corrupted DNA-binding specificity and ectopic transcription underpin dominant neomorphic mutations in KLF/SP transcription factors. BMC Genomics 20, 417

66. Mitchell WB G. M., Jaffray JA, et al. (2011) Case Report of Erythroid Transcription Factor EKLF Mutation Causing a Rare Form of Congenital Dyserythropoetic Anemia in a Patient of Taiwanese Origin. in ASH Annual Meeting Abstracts

67. Helias, V., Saison, C., Peyrard, T., Vera, E., Prehu, C., Cartron, J. P., and Arnaud, L. (2013) Molecular analysis of the rare in(Lu) blood type: toward decoding the phenotypic outcome of haploinsufficiency for the transcription factor KLF1. Hum Mutat 34, 221–228

68. Soni, S., Pchelintsev, N., Adams, P. D., and Bieker, J. J. (2014) Transcription factor EKLF (KLF1) recruitment of the histone chaperone HIRA is essential for beta-globin gene expression. Proc Natl Acad Sci U S A 111, 13337–13342

69. Sengupta, T., Chen, K., Milot, E., and Bieker, J. J. (2008) Acetylation of EKLF is essential for epigenetic modification and transcriptional activation of the beta-globin locus. Mol Cell Biol 28, 6160–6170

70. Brown, R. C., Pattison, S., van Ree, J., Coghill, E., Perkins, A., Jane, S. M., and Cunningham, J. M. (2002) Distinct domains of erythroid Kruppel-like factor modulate chromatin remodeling and transactivation at the endogenous beta-globin gene promoter. Mol Cell Biol 22, 161–170

71. Chen, X., and Bieker, J. J. (2004) Stage-specific repression by the EKLF transcriptional activator. Mol Cell Biol 24, 10416–10424

72. Quadrini, K. J., and Bieker, J. J. (2002) Kruppel-like zinc fingers bind to nuclear import proteins and are required for efficient nuclear localization of erythroid Kruppel-like factor. J Biol Chem 277, 32243–32252

73. Sengupta, T., Cohet, N., Morle, F., and Bieker, J. J. (2009) Distinct modes of gene regulation by a cell-specific transcriptional activator. Proc Natl Acad Sci U S A 106, 4213–4218

74. Southwood, C. M., Downs, K. M., and Bieker, J. J. (1996) Erythroid Kruppel-like factor exhibits an early and sequentially localized pattern of expression during mammalian erythroid ontogeny. Dev Dyn 206, 248–259

75. Vorackova, I., Suchanova, S., Ulbrich, P., Diehl, W. E., and Ruml, T. (2011) Purification of proteins containing zinc finger domains using immobilized metal ion affinity chromatography. Protein Expr Purif 79, 88–95

76. Bailey, T. L., Boden, M., Buske, F. A., Frith, M., Grant, C. E., Clementi, L., Ren, J., Li, W. W., and Noble, W. S. (2009) MEME SUITE: tools for motif discovery and searching. Nucleic Acids Res 37, W202–208

